# Causality Mapping Using Resting-state fMRI Reveals Hyperactivity and Hypoconnectivity in Schizophrenia Patients

**DOI:** 10.1101/2024.09.25.614909

**Authors:** Alishba Tahir, Wajiha Abdullah, Wjeeh ul Azeem, Muhammad Farhan Khalid, Turki Abualait, Sadia Shakil, Fayyaz Ahmed, Shahid Bashir, Safee Ullah Chaudhary

## Abstract

Schizophrenia (SZ) is a debilitating disorder in which patients exhibit psychotic behavior due to aberrant connectivity between different regions of the brain. Advances in neuroimaging have now enabled the diagnosis and analysis of SZ in order to elucidate the whole brain functional connectivity networks. In the present study, we have used resting-state functional magnetic resonance imaging (rs-fMRI) to elucidate the causal relationships amongst the differentially activated brain regions between SZ patients (n=10) and healthy controls (n=10). Vector auto-regression (VAR) model and Granger causality (GC) were then applied to construct a functional connectivity network and analyze the causal effects in SZ patients. Our results revealed that the average voxel activation in the frontal lobe (FL), basal ganglia (BG), and ventricular system (VS) was significantly higher in patients indicating hyper-activity as compared to controls. Conversely, cerebellum white matter (CBWM) showed higher activation in the controls as compared to patients. A higher Pearson correlation was observed between the controls as compared to patients while VAR and GC showed higher functional connectivity among all the regions of interest (ROIs) along with more causal relations in the controls. Finally, mediation analysis showed that right middle superior frontal gyrus acts as a strong partial mediator between left accumbens area and left middle superior frontal gyrus. Taken together, this study decodes the dysregulated brain activity in schizophrenia showing hyperactivation in patients when compared with the healthy controls which leads to alterations in neural connections resulting in hypoconnectivity.

## 1 Introduction

Schizophrenia (SZ) is a chronic mental disorder that affects over 24 million people across the globe (*Schizophrenia*, no date). The disease onset may begin during late adolescence and has a lifetime prevalence of 2.3% (*The epidemiology of early schizophrenia. Influence of age and gender on onset and early course - PubMed*, no date; Gogtay *et al*., 2011; Subramaniam *et al*., 2021). SZ-induced health deterioration is known to lower patients’ life expectancy by 15-25 years besides degrading the overall quality of life (Wildgust, Hodgson and Beary, 2010; Vos *et al*., 2017). SZ is divided into three types based on its clinical symptoms; 1. Positive symptoms including hallucinations, delusions, and disorganized behavior (*Psychiatry*.*org - DSM-5 Fact Sheets*, no date; Buckley and Miller, 2015; Bighelli *et al*., 2018), 2. negative symptoms, such as flat affect and lack of movement etc. (Hare *et al*., 2019), and 3. cognitive symptoms, such as disorganized thinking, confusion and memory loss (Hare *et al*., 2019). Molecular investigations into the mechanistic underpinnings of SZ have led to discovery of several genetic alterations such as single-nucleotide polymorphisms and copy number variations in genes regulating dopaminergic neurotransmission in the brain (Li *et al*., 2011; Warburton *et al*., 2016).These molecular changes in SZ result in significant functional alterations in the neuronal connections between various regions of the brain and cause functional dysregulation (Konradi and Heckers, 2003; Lodge and Grace, 2011a; Beveridge and Cairns, 2012). In particular, altered patterns of neural activation in the prefrontal cortices, including the anterior cingulate cortex (ACC), the supplementary motor area (SMA), the pre-SMA, the parietal cortex, and the subcortical basal ganglia nuclei (Huang *et al*., 2018; Hua *et al*., 2020; Zhang *et al*., 2020) are known indicators of SZ.

Chronic SZ causes rewiring of the brain thus altering the connections between different regions. Bioimaging research into SZ has reported a significant reduction in the functional connectivity (FC) amongst various regions of the brain(Koenig *et al*., 2001; Skudlarski *et al*., 2010; Zhou *et al*., 2010; Erdeniz *et al*., 2017; Dong *et al*., 2018). To decode the role of FC in SZ, researchers have studied the individual regions of the brain and evaluated their interrelationships towards uncovering the underpinning mechanisms. Individuals suffering from SZ are known to exhibit aberrations in regional connectivity which is caused by hyper/hypo activation in various regions of the brain ^27–30^. The affected areas include all the major lobes of the brain including the frontal lobe (FL), temporal lobe (TL), occipital lobe and the memory center, the hippocampus (HC). FL which is known to be involved in cognition, memory, and higher order functions (Chayer and Freedman, 2001; Gillingham *et al*., 2017), escalates the progression of SZ by lowering its activity (Corponi *et al*., 2021) and impairing connectivity with the other regions of the brain (Jiang *et al*., 2018). This results in induction of hallucinations and delusions which are two of the most prevalent symptoms observed in SZ patients (Jiang *et al*., 2018). Shinba *et al*. and Chatterjee *et al*. investigated the abnormal functionality of FL in SZ and measured its activity through near-infrared spectroscopy (NIRS) (Shinba *et al*., 2004) and fMRI (Chatterjee *et al*., 2020), respectively. Such structural and functional dysfunction is not only limited to FL but also reported in other regions of the brain. For instance, OL which constitutes the visual cortex of the brain (Schneider *et al*., 2019), has been reported to have reductions in the grey matter volume in the lateral and medial regions during SZ (Jiang *et al*., 2018; Tordesillas-Gutierrez *et al*., 2018). It is important to note that this atrophy manifests without any deterioration in the visual capacity of the patients (Johnson *et al*., 2015). Similarly, TL is also affected during the progression of SZ; Spalthoff *et al*. reported altered gyrification in parts of TL including bilateral insula and temporal pole along with decreased grey matter volumes (Spalthoff, Gaser and Nenadić, 2018). HC, a part of the medial TL, is a key operating brain structure that is involved in memory retention and retrieval (Squire, Stark and Clark, 2004) (Huijgen and Samson, 2015) and its dysfunction furthers the cognitive symptoms related to SZ (Ghoshal and Conn, 2015). Note that HC plays a role in healthy and diseased brain as the dysfunction following SZ causes hippocampal dysregulation which leads to dopaminergic dysfunction (Lodge and Grace, 2007, 2011b; Grace, 2012).

Interestingly, in complete contrast with these findings, Salisbury *et al*. found higher resting state cerebral blood flow (rsCBF) in the bilateral temporal lobe and putamen indicating aberrant hyperactivity (Salisbury *et al*., 2022). Zovetti *et al*. performed a task-based fMRI experiment and found out that patients’ white matter tracts were hyperactivated inducing hyperconnectivity in areas encompassing the corpus callosum (Zovetti *et al*., 2022). A meta-analysis by Wu *et al*. revealed multiple over-activated areas including bilateral dorsolateral prefrontal cortex, posterior parietal cortex, anterior cingulate cortex, anterior insula and supplementary motor area - areas making up the salience and central executive network (Wu and Jiang, 2020). Johnsen et al. tested brain activation in blood relatives of SZ patients through task-based fMRI in children and adolescents and found hyperactivation in the right dorsolateral prefrontal cortex and left caudate head, the left inferior frontal cortex, bilateral caudate, left inferior temporal gyrus, and bilateral frontal pole (Johnsen *et al*., 2020). This hyperactivity was used to construct connectivity maps showing aberrant connectivity in SZ patients. Salvador et al., found over-connected regions with the default mode network (DMN) which exhibited the most hyperactivity. Other areas that have been also found to be over connected in brain disorders include several cortical and sub-cortical structures (Salvador *et al*., 2010a). To exemplify, Koshiyama et al., reported over-connected brain networks between the major brain areas including the frontal, temporal and occipital lobe (Koshiyama *et al*., 2020).

Researchers have also endeavored to assess white matter composition and identified aberrant activations in several key brain regions, particularly the frontal lobe (FL), which is involved in the facilitation of advanced cognitive processes (Brewer *et al*., 2007). Brewer et al. conducted a longitudinal PET study and found an increased activation of the FL in SZ patients which could attribute towards the positive symptoms of SZ (Brewer *et al*., 2007). Similarly, basal ganglia (BG) which are present deep within the brain are also reported to be hypoactive and hyperactive as the disease progresses (Salvador *et al*., 2010b). Changes in size and structure of BG have been reported in majority of the cases thus making it one of the key players in SZ. The ventricular system (VS), located near the BG, has shown larger volumes in SZ patients which can lead to a hyperactive state (Algunaid *et al*., 2018). Another close by area is the cerebellum white matter (CWM) which is responsible for executive control and motor movement hence important from the point of view of psychopathology (Hariri, 2019). Cerebellum white matter (CWM) has been reported to increase in size contributing to the positive symptoms of SZ but with continued treatment it decreases over time (Christensen, Holcomb and Garver, 2004).

Till date, the distribution and variation in FC affecting neuronal communication and its underlying pathophysiological mechanisms, remains unclear (Rubinov and Bullmore, 2013). However, recent spatiotemporal investigations of the underlying neuropathological mechanisms of the diseased brain have provided some new insights (Minzenberg et al., no date; Snitz et al., 2005; Yoon et al., 2008; Lynall et al., 2010; Rubinov and Bullmore, 2013). In particular, studies leveraging technologies such as magnetic resonance imaging (MRI) (Sutcliffe et al., 2016; Vanes et al., 2019) and functional magnetic resonance imaging fMRI (del Re et al., 2019) have been employed to report a wide variety of abnormal structural and functional connections among anatomically distant regions of the brain in SZ patients (Sutcliffe et al., 2016; del Re et al., 2019; Vanes et al., 2019). Ralf et. al., employed a combination of fMRI and diffusion tensor imaging (DTI), and reported a modified structure-function relationship in SZ (Chayer and Freedman, 2001). Liang et al. further decoded the temporal associations between various regions of the brain and analyzed the FC by calculating multiple neuroimaging characteristics such as gray matter volume (Gillingham et al., 2017).

Towards investigating FC in SZ using rsfMRI, we have elucidated the causal relations between ROIs. For that, spatial independent component analysis (ICA) (Yang et al., 2020) was used to identify the significant features in fMRI followed by computing voxel activations for the whole brain. The results showed differential activations for 16 areas in patients which were used for further statistical analyses. A seed-based temporal correlation analysis (SCA) (Mannell et al., 2010) was used to assess time-series connections among the ROIs. Resting-state functional connectivity (rsFC) and causal interactions among the regions were evaluated through multivariate vector autoregression (VAR) (Granger, 1969) and Granger causality (GC)(Wismüller and Vosoughi, 2021). Our results from spatial ICA indicated a higher BOLD signal in FL, BG, VS and CWM in SZ patients. Pearson correlation showed more significant correlations in all ROIs in controls compared to the patients. Significantly increased functional connections in vector auto regression were also observed amongst the controls along with increased G-causal relations between ROIs. Lastly, mediation analysis revealed that right middle frontal superior gyrus acts as a partial mediator between left accumbens and left middle frontal gyrus in both patients and control cases with a stronger partial mediation in patients.

In conclusion, this study provides novel evidence of hyperactivity in all ROIs in SZ patients along with hypoconnectivity with decreased causal connections as the disease progresses. These causal inferences between the four selected regions of the brain expand our understanding of the resting-state FC (rsFC) in patients with SZ at the whole-brain level. This study also helps to characterize the FC among different regions of the brain and its variations in patients with SZ thereby providing valuable insights into the neural basis of SZ. Taken together, the study has the potential to direct SZ diagnosis through the identification of neuroimaging markers, decoding the functional connectivity networks and elucidating the causal interactions in the healthy as well as the diseased brain (Abboud, Noronha and Diwadkar, 2017), (Acar et al., 2019; Fan et al., 2021).

## 2 Methods

### 2.1 Participants

The study was approved by the local ethic committee of King Fahad Specialist Hospital Dammam (KFSHD) (NEU0334). A total of 20 male participants were selected (Age; 33.14 ± 9.96, mean ± SD), 10 of whom were healthy controls, and the others were schizophrenia (SZ) patients. Individuals from the healthy cohort were randomly chosen whereas the SZ patients were selected from home-grown psychiatric clinics and were clinically stable for two weeks prior to testing. The patients were previously diagnosed based on the DSM-IV (Bell, 1994) criteria followed by manual confirmation from an experienced research assistant. All participants gave written informed consent for participation. Participants were excluded if they: (a) met DSM-IV criteria for substance dependence or severe/moderate abuse during the prior six months; (b) had a clinically unstable or severe general medical disorder; or (c) had a history of head injury with documented neurological sequelae or loss of consciousness. Psychopathology was assessed by trained research assistants using the scale for the assessment of negative symptoms (SANS) and the scale for the assessment of positive symptoms (SAPS)(Andreasen *et al*., 1995). Specific subscale scores were summed to derive measures of positive symptoms (hallucination and delusion subscales), disorganization (formal thought disorder, bizarre behavior, and attention subscales), and negative symptoms (flat affect, alogia, anhedonia and demotivation subscales).

### 2.2 Resting state functional magnetic resonance imaging

A Siemens Magnetom Verio 3T MRI clinical scanner (Siemens Healthineers) and a 12-channel phased-array head coil were used to acquire: (1) T1-weighted 3D magnetization-prepared rapid gradient-echo imaging (MPRAGE): Repetition Time (TR) = 1600 ms, Echo Time (TE) = 2.19 ms, inversion time = 900 ms, flip angle = 9°, acquisition plane = sagittal, voxel size = 1 × 1 × 1 mm^3^, field of view (FOV) = 256 mm, acquired matrix = 256 × 256, acceleration factor (iPAT) = 2; (2) Fluid attenuated inversion recovery (FLAIR): TR = 9000 ms, TE = 128 ms, inversion time = 2500 ms, flip angle = 150°, acquisition plane = axial, slice thickness = 5mm, FOV = 220 mm, acquired matrix = 256 × 196, acceleration factor (iPAT) = 2. A total of 404 scans were taken per subject and the first four scans were discarded before performing the pre-processing which was carried out using statistical parametric mapping 12 (SPM 12).

To reduce the head movement in scans, each scan was corrected for motion against the mean image and realigned. Next, the scans were processed for spatial smoothing by taking the voxel averages. Gaussian kernel was used for images which had a normal distribution form. After spatial smoothing, the images were normalized according to the template image EPII (Echo-planar imaging) and specified to level one by using the binary condition and specified at level 2 by using all fifteen contrasts to undertake a group analysis.

### 2.3 Statistical Data Analysis

Mann Whitney test was used to compare the voxel activations between the controls and patients. To check for a correlation among the ROIs in both the groups, Pearson correlation was computed in MATLAB. To check if there were any time lagged causal connections amongst the ROIs vector auto-regression was computed and these results were further validated by Granger causality (GC). Spatial independent component analysis (ICA) was used to elucidate the connectivity patterns between all the brain areas in SPM towards determining their causal relationships (Gohel *et al*., 2018). Seed based correlation analysis (SCA) was employed to evaluate the correlation between “seeds” (clusters of neurons) that show concurrent activity in the 16 regions of interest (ROIs) that were differentially activated in patients compared to the controls(Beckmann *et al*., 2005; Calhoun and Adali, 2012; Sheffield and Barch, 2016; Calhoun and de Lacy, 2017). Finally, vector auto-regression (VAR) and Granger causality (GC) were performed to study the bidirectional interaction amongst the ROIs and the time-lagged causal effects on each ROI, respectively(Zhou *et al*., 2009; Guo, Liu, Liu, *et al*., 2015; Silverstein, Bressler and Diwadkar, 2016).

### 2.4 Estimation of Granger Causality

GC(Seth, Barrett and Barnett, 2015) was modelled using equations 1 and 2 to analyze the fMRI data and elicit the causal connectivity between the activated regions of the brain. Specifically, in conventional regression models, any change in endogenous variable X_*t*_ results in a change in exogenous variable Y_*t*_ - that is, X_*t*_ causes Y_*t*_ and not vice versa (**Figure 1A**). To incorporate a bi-directional causality, vector auto-regression (VAR) models treat all variables equally such that X_*t*_ can cause Y_*t*_ (as well as their lags), and vice versa (**Figure 1B & C**). The VAR models for two of the four regions of interest (ROIs) i.e. FL and TL are defined in equations 1 and 2, respectively:

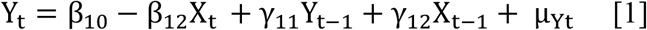

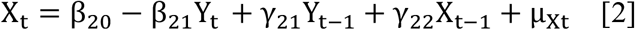

**Figure 1.**
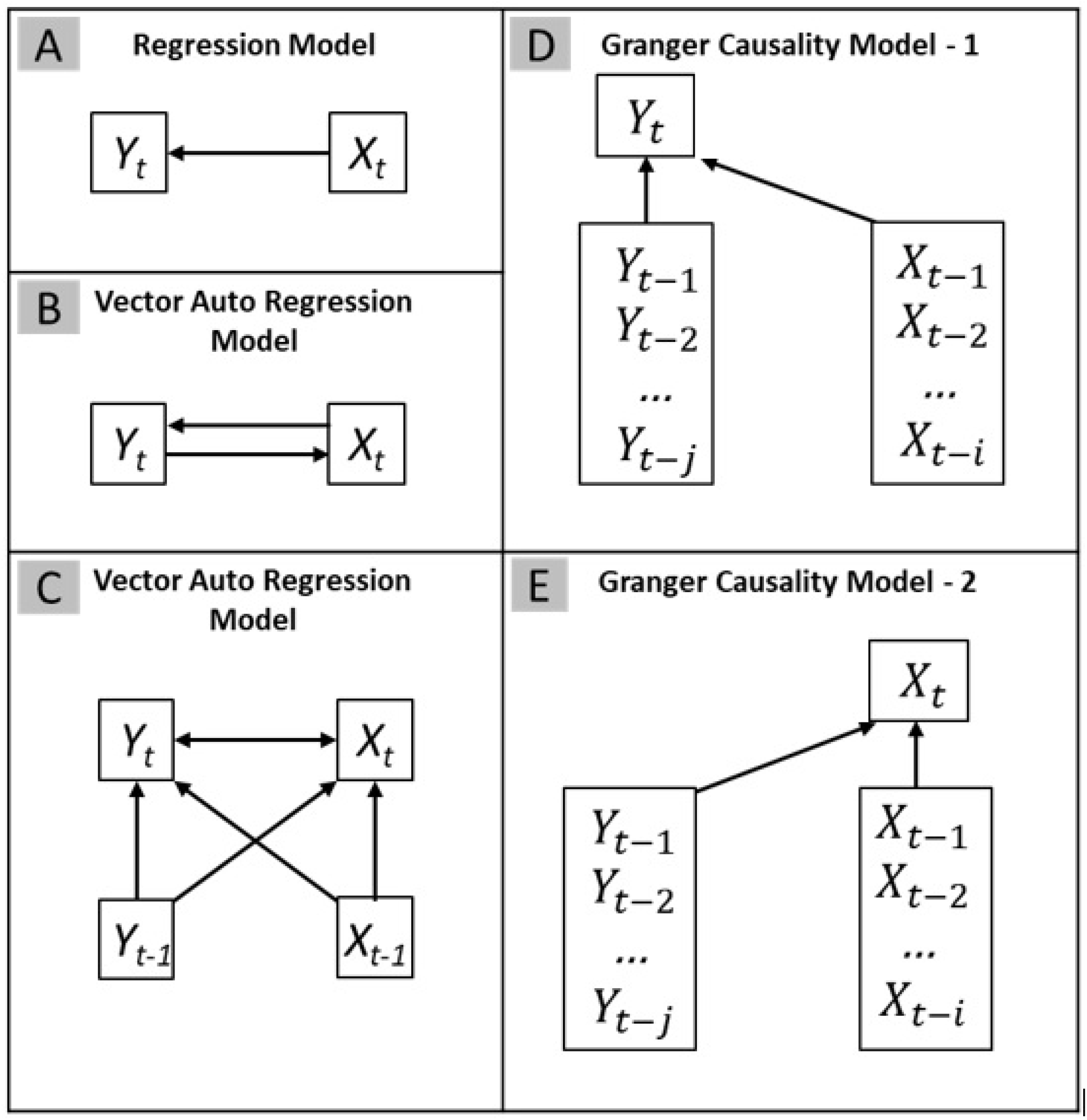
Causal relations between variables. (A) Regression Model: *X*_*t*_ causes *Y*_*t*_, (B) Vector Auto-regression Model: *X*_*t*_ can cause *Y*_*t*_ and vice versa, (C) Vector Auto-regression Model: *X*_*t*_ can cause *Y*_*t*_ and their lags, and vice versa, (D) Granger Causality Model-1: *Y*_*t*_ may be affected by changes in lag values at *X*_*t-1*_, *X*_*t-2*_ and *X*_*t-i…*_, and changes in lag values in *Y*_*t-1*_, *Y*_*t-2*_ *and Y*_*t-j*_,.. where i and j represent different time points (E) Granger Causality Model-2: X_*t*_ may be affected by changes in lag values at *Xt*_*-1*_, *X*_*t-2*_, *X*_*t-i*_*…* and changes in lag values in *Y*_*t-1*_, *Y*_*t-2*_, *Y*_*t-i*_*…*

Where *Y*_*t*_ and *X*_*t*_ are stationary, *µ*_*Xt*_ and *µ*_*Yt*_ are not correlated and the resultant models are first order VAR models (Dimitrios Asteriou, no date).

Model [1] and [2] can be re-written as:

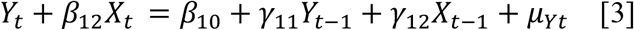

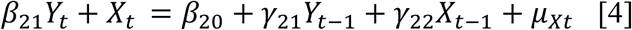

Or in the matrix form as (Arnold, Liu and Abe, 2007):

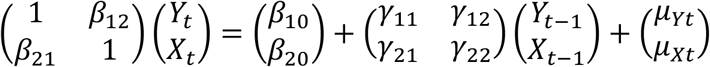

And in the short form as:

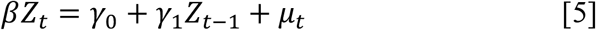

Pre-multiplication with *β*^−1^gives:

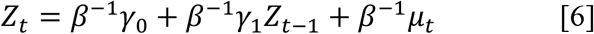

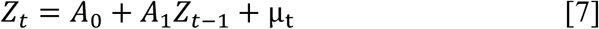

Model [7], the reduced form of bivariate auto regression (BVAR) model, with all variables considered as stationary is written as *A* = (Â_0_,Â_1_)^*T*^ by ordinary least squares (OLS) and the equation as:

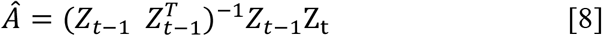

Where 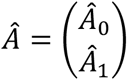, and 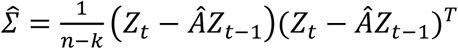 and *k* = number of parameters.s

**Figure 1D and E** show that *Y*_*t*_ may be affected by changes in lag values at *X*_*t-1*_, *X*_*t-2*_ and *Xt*_*-I*_, *Y*_*t-1*_, *Y*_*t-2*_ *and Y*_*t-I*_ and X _*t*_ may be effected by changes in lag values at *Xt*_*-1*_, *X*_*t-2*_, *X*_*t-i*,_ *Y*_*t-1*_, *Y*_*t-2*_ and *Y*_*t-i*_

These GC models provide the following equations(Seth, Barrett and Barnett, 2015):

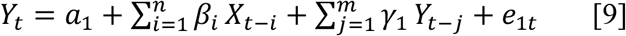

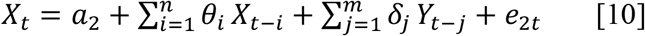

The GC models estimated, using Python for equations [9] and [10], provided four possible parameters i.e.(i) the lagged X terms in equation [9] may be statistically different from zero as a group, (ii) the lagged Y terms in equation [10] may be statistically different from zero as a group, and the lagged Y terms in equation [1] may not be statistically different from zero, hence Y causes X, (iii) Both X and Y are statistically different from zero in [9] and [10], hence both X and Y cause each other. (iv) Both sets of X and Y terms are statistically not different from zero in [9] and [10], therefore both X and Y do not cause each other.

Following procedure was applied to equations [9] and [10] to calculate the G-causal relations amongst the ROIs:

**Step 1**: Regressing *Y*_*t*_ on lagged term *Y*_*t™j*_ *as*:

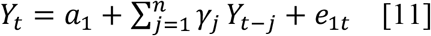

**Step 2**: Regressing *Y*_*t*_ on lagged terms of both, *Y* and *X*:

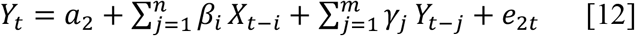

OLS models were used on equations [4] and [5] to calculate an unrestricted residual sum of square.

**Step 3**: Testing the effects on constant variables:

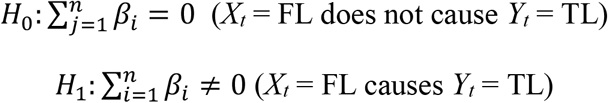

**Step 4**: calculating F-statistic:

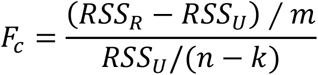

where m = lags length at Y and k = number of parameters, and k = m + n + 1

Step 5: Applying Bayesian information criterion (BIC)

Lags length (*m*) was computed by the Bayesian Information Criterion (BIC) to compare models:

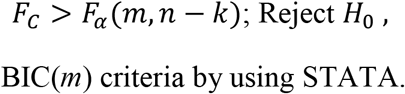

This resulting novel GC model was applied to the results of the VAR model to establish the causal links between ROIs.

## 3 Results

### 3.1 Frontal lobe, basal ganglia, ventricular system and cerebellum white matter show hyperactivity in patients with Schizophrenia

To evaluate the differences between brain activation patterns in schizophrenic patients, resting state functional magnetic resonance imaging (rsfMRI) scans of 10 healthy controls (**Supplementary Data 1**) and 10 SZ patients (**Supplementary Data 2**) were obtained for 400 seconds. The recorded scans were pre-processed for realignment, spatial normalization, smoothing, and co-registration using statistical parametric mapping 12 (SPM12)(*SPM12 Software - Statistical Parametric Mapping*, no date). The voxel activations were deduced using the blood oxygen level dependent (BOLD) signal in the sixteen regions of interest (ROIs) which have been majorly anatomically categorized into frontal lobe (FL), basal ganglia (BG), ventricular system (VS), and cerebellum (CB). The voxel activations were then compared between healthy controls and SZ patients (**Figure 2**). Mann Whitney analysis showed significant differences in the average brain activity in the two groups for all ROIs. Right lateral ventricle, 3^rd^ ventricle, left superior frontal gyrus and right superior frontal gyrus show the highest voxel activations in both groups, while right triangular part of the inferior frontal gyrus show the least activation in controls as well as the patients. Right cerebellum white matter is the only area that is more active in the controls as compared to the patients. Hence, patients exhibited an overall higher activity in all ROIs except one region, suggesting a hyperactive brain state.

**Figure 2.**
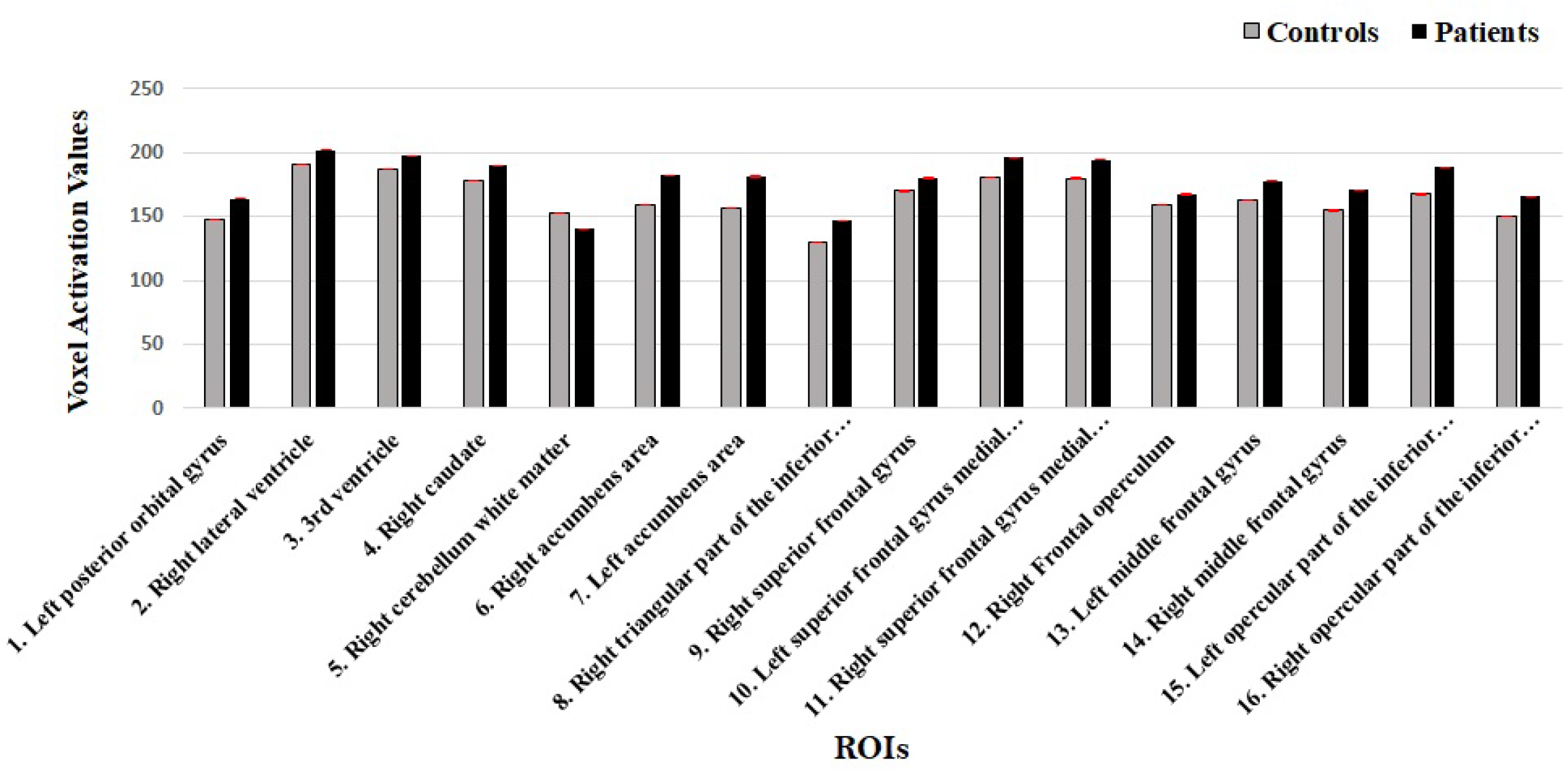
Average Voxel Activation in Controls and Schizophrenia Patients. The bar graph shows average voxel activations in 16 ROIs for controls (grey) and patients (black).

### 3.2 Simultaneous activation patterns in ROIs differ significantly in Schizophrenia

To evaluate whether the aberrant activations observed within each ROI altered patterns of simultaneous activations, Pearson correlation was computed between ROIs for both controls and patients. In the VS, the right lateral ventricle (RLV) had the highest correlation with the right caudate (r = 0.84384, p < 0.05) of BG. In FL, the left posterior gyrus (LPG) showed the greatest correlation with the left accumbens area (LA) in BG. The inverse relationship was also the strongest amongst BG, as the left accumbens area (LA) had the strongest correlation with the left posterior orbital gyrus (LPOG) in the FL (r = 0.90321, p < 0.05). In the CWM, the strongest correlation was with the Right Superior Frontal Gyrus in the Frontal Lobe (r = 0.68393, p < 0.05). In the controls the right lateral ventricle sub region (RLVSR) of the VS exhibited the strongest correlation with the right caudate in the BG (r = 0.85632, p < 0.05). The FL had the strongest correlation with the right accumbens area (RA) of BG through the right superior frontal gyrus (RSFG) (r = 0.74415, p < 0.05). The right caudate in the BG had the greatest correlation with the right lateral ventricle (RLV) in VS (r = 0.85632, p < 0.05) and RCWM in the CWM had the greatest correlation with the right accumbens (RA) in the BG (r = 0.65585, p < 0.05). The results showed 222 significant correlations for SZ patients amongst all ROIs and 232 significant correlations in the controls thus indicating an altered ROI connectivity in SZ patients **(Figure 3)**.

**Figure 3.**
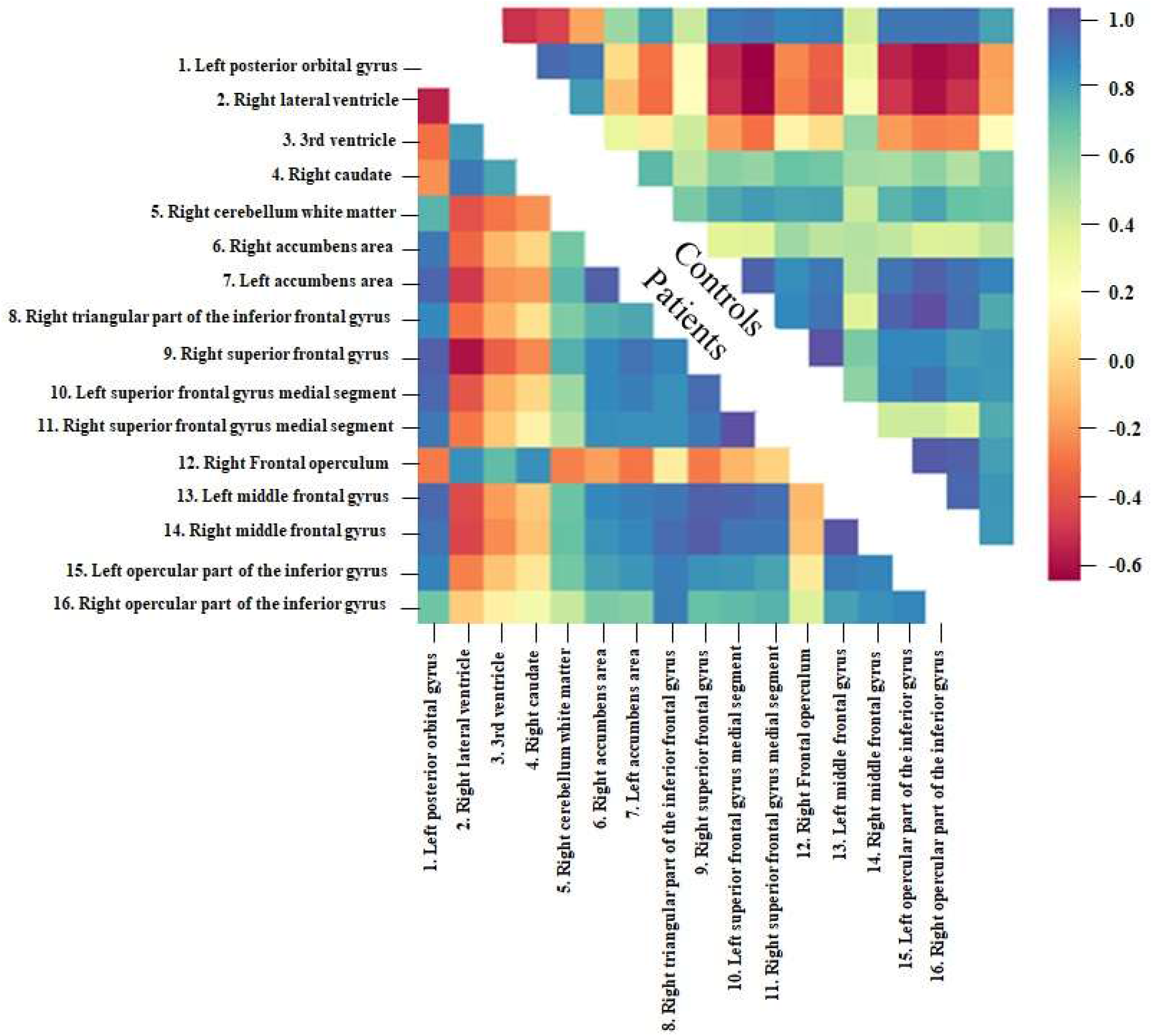
Correlation Between ROIs in Controls and Schizophrenia Patients. The heat map shows Person correlation between the ROIs in controls and SZ patients.

### 3.3 Schizophrenia patients show hyper-connectivity between the ROIs

To examine the effect of altered correlation between ROIs, the temporal variations in functional connectivity (FC) between the ROIs were evaluated (Venkataraman *et al*., 2012) at times *t*-1 and *t*-2 using vector auto-regression (VAR) model (Chen *et al*., 2011), pair-wise, for all ROIs. Lags *t*-1 and *t*-2 were calculated through autocorrelation which showed connectivity with the other ROIs until *t*-2 seconds. In controls **(Figure 4A)**, FL exhibited strong causal connectivity between the right superior frontal gyrus (Right MSFG) and the right cerebellum white matter (Right CWM) (t-2= 0.236, p < 0.05) whereas in the patients **(Figure 4B)**, it was between the left superior frontal gyrus medial segment (left MSFG) and the Right CWM. For controls, BG had significant connectivity with the Right CWM (t-1= 0.1820448, p < 0.05), whereas in patients’ BG showed significant connectivity with the right lateral ventricle in VS (t2 = 2103896, p<0.05). In controls, VS showed connectivity with the left accumbens (Left acc) in the BG (t-2= 0.1689018, p < 0.05) while in the patients, the ^3rd^ ventricle had significant connectivity with the Right MSFG in the FL (t-1= 0.4509556, p < 0.05). Overall, controls had 41 significant connections between ROIs whereas patients had 38 indicating that functional connectivity is reduced in SZ. Hyperactivity in the patients did not accompany hyperconnectivity; the reduced connectivity led to hyperactivation in the ROI’s. To validate the structural model of the brain connectivity network obtained through VAR and examine the intercausal effects of the ROIs in the controls and patients, Granger causality test was employed. In controls (**Figure 5A**) right lateral ventricle (RLV) G-caused right superior frontal gyrus (RSFG) (chi-square=35.337) in VS and G-caused right triangular inferior part of the frontal gyrus (RTriFG). RSFG G-caused third ventricle (chi-square=11.724) from the VS and RTriFG also G-caused right accumbens. Right caudate G-caused right frontal operculum (chi-square=20.462) in BG and right accumbens G-caused left opercular part of the inferior frontal gyrus (Left OpIFG). In patients (**Figure 5B**) RLV G-caused right accumbens (chi-square=20.748) in VS and right lateral ventricle also G-caused left OpIFG. RSFG G-caused right cerebellum white matter (chi-square=24.611) in FL and left superior frontal gyrus medial segment (LMSFG) G-caused right accumbens. Right caudate G-caused RSFG (chi-square=22.809) and right caudate also G-caused RTriFG. Overall, the controls had a total of 67 significant G-causal relationships compared to 52 in the patients confirming the results of the VAR model. The results of GC also indicate that loss in connections has a more magnified effect on the loss in useful causal connections as GC shows a more steep decrease in the causal interactions amongst the patients.

**Figure 4.**
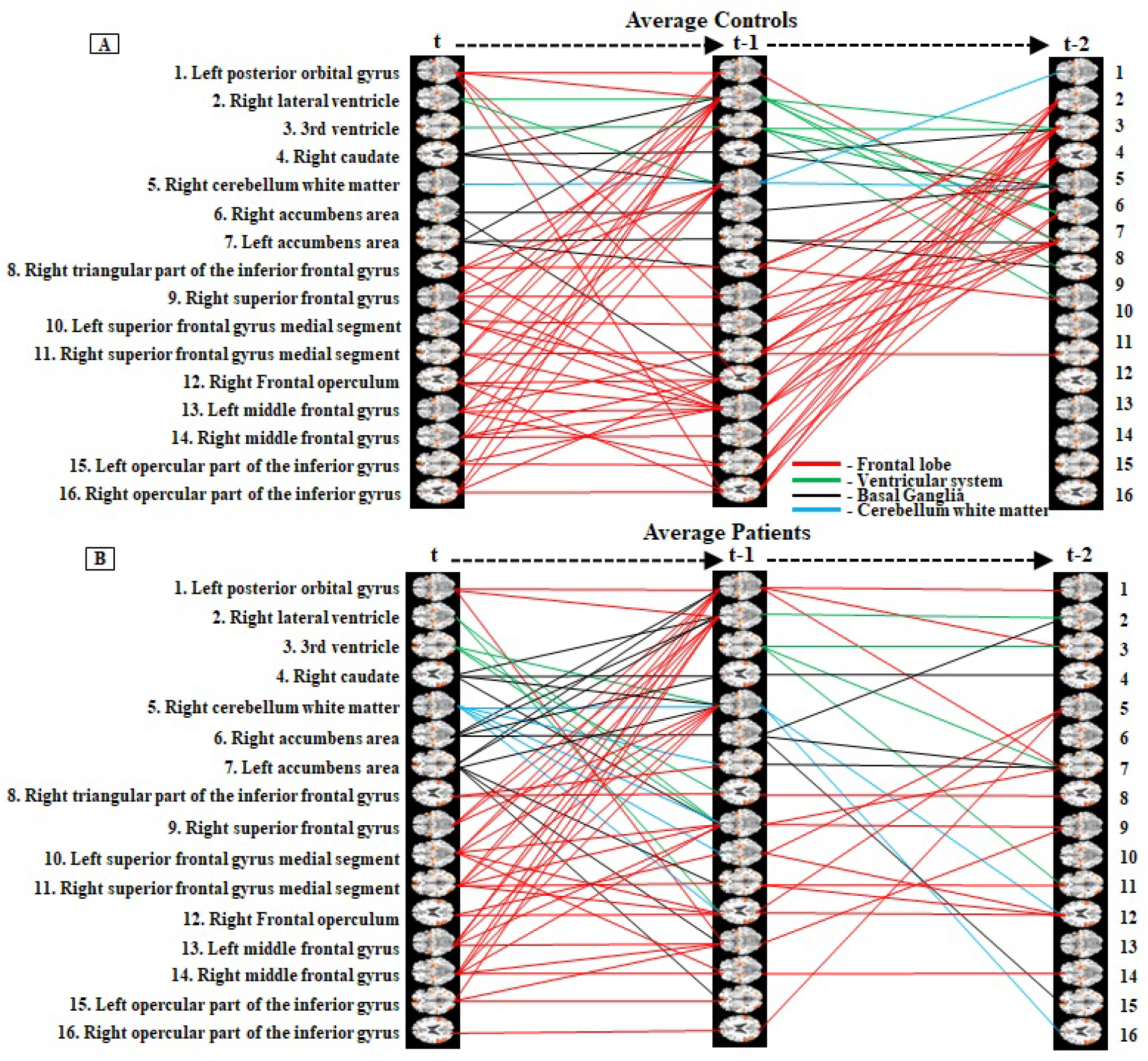
Structural Connectivity in ROIs through vector autoregression model. Significant causality links between the ROIs in the patients (A) and the controls (B).

**Figure 5.**
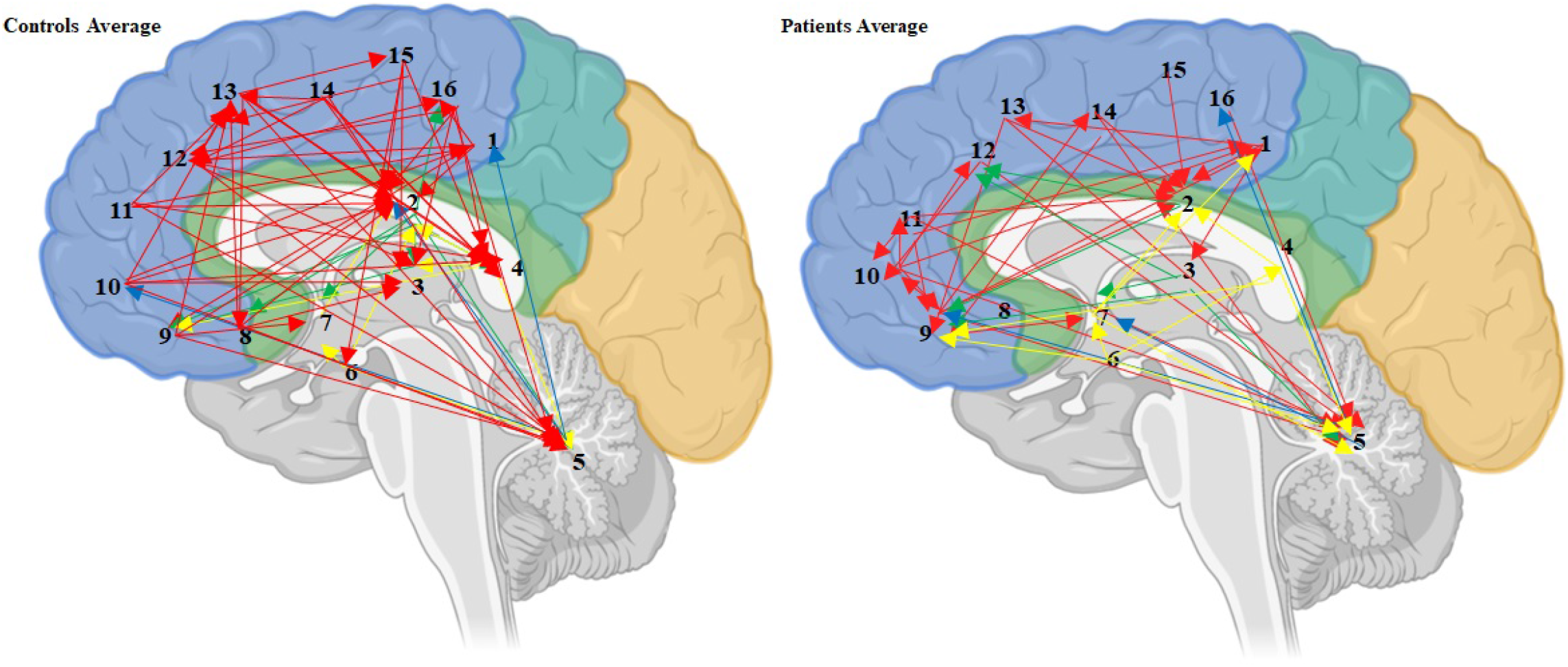
Structural Connectivity in ROIs obtained through Granger Causality. Arrows show significant granger causality between ROIs for the controls (A) and patients (B).

### 3.4 Right middle superior frontal gyrus acts as a partial mediator between left accumbens and left middle superior frontal gyrus

To investigate whether one of the ROIs mediates the relationship between the other ROIs, all possible combinations with 16 ROIs for mediation analyses were tested. Highest partial mediation was observed between nucleus accumbens (NA) (independent variable) and left middle superior frontal gyrus (LMSFG) (dependent variable) with right middle superior frontal gyrus (RMFSG) as a mediator for both controls and SZ patients (Figure 6A and B). However, in controls, RMSFG shows relatively lower mediation with the value of -0.0398 (p<0.05) whereas patients exhibited a stronger partial mediation value of 0.5704 (p<0.05) **(Figure 6)**. These results are indicative of an over-compensation in the patients as the regions of interest show a higher effect of removing one brain region in patients as compared to the controls. These results reconfirm that aberrant activation patterns lead to loss in functional connectivity.

**Figure 6.**
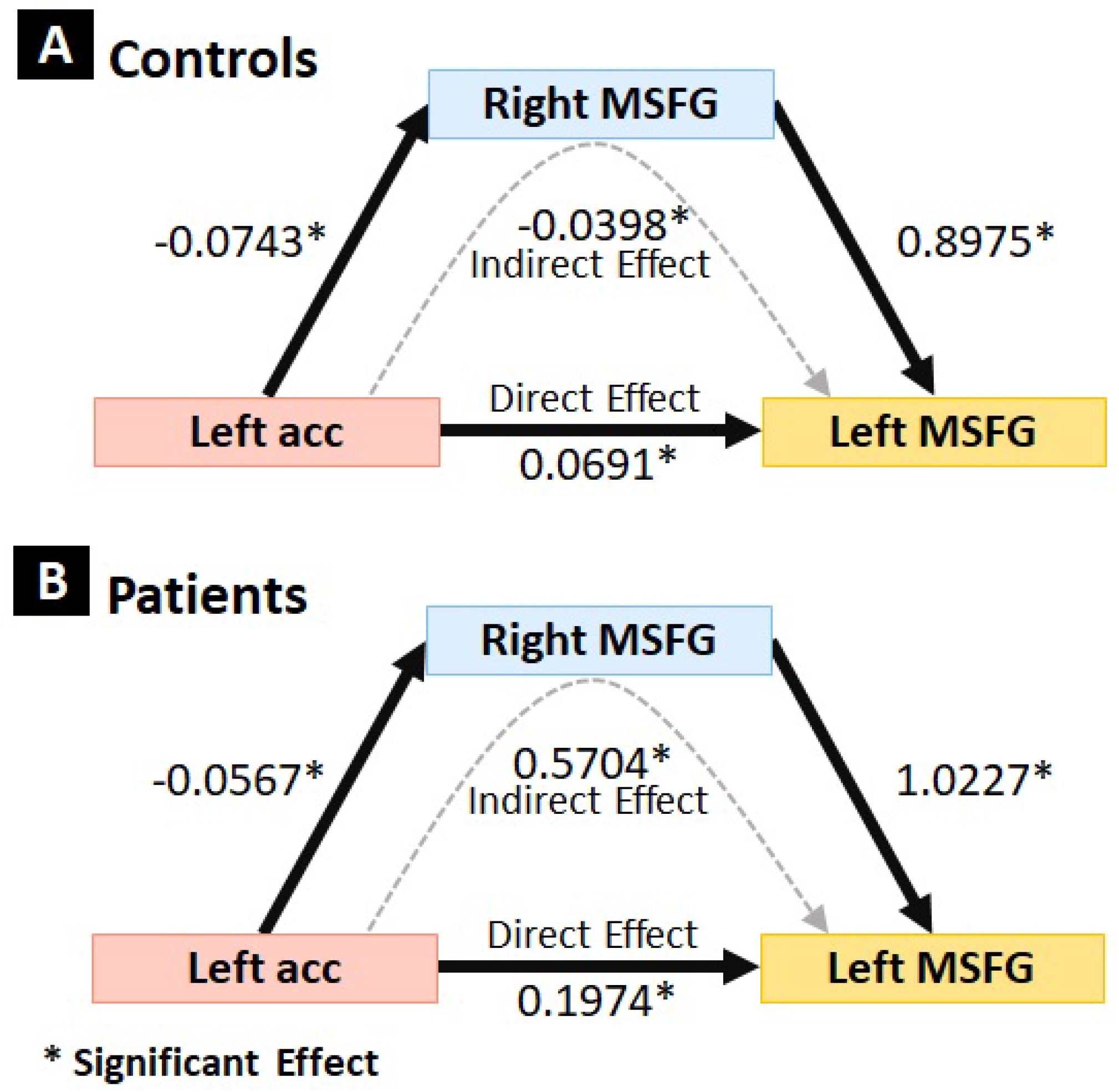
Mediation Analysis. **(A)** Mediation in controls between Left accumbens area, right MSFG, and left MSFG, **(B)** Mediation in patients between Left accumbens area, right MSFG, and left MSFG.

## 4 Discussion

In this study, we aimed to establish the resting state functional connectivity (rsFC) patterns in healthy controls and schizophrenia (SZ) patients. Functional magnetic resonance imaging (fMRI) was used to detect the regional cerebral blood flow (rCBF) to calculate the average voxel activations for the controls as well as the patients. The data was then analyzed for computing the causal relations between ROIs using vector auto-regression (VAR) and Granger causality (GC).

Our results showed an increase in the voxel activations in the FL (frontal lobe), BG (basal ganglia), VS (ventricular system) and CB (cerebellum) in SZ patients suggesting hyperactivity in this patient cohort. Previously, both high and low voxel activations have been reported in SZ patients (Bobilev, Perez and Tamminga, 2020)(Holt *et al*., 2006). Positron emission tomography (PET) scans show higher blood flow in the dentate gyrus (DG; a part of the HC) which receives sensory input from the brain to create memories (Pattabiraman *et al*., 2021). The positive symptoms (hallucinations and delusions) of SZ could also be a result of this altered activity. Frontal lobe (FL) is one of the areas that exhibits higher activation in SZ patients. The left posterior orbital gyrus (POG) exhibits hyper-activation in SZ patients and this contributes towards the symptoms of psychosis. While the literature does not specifically mention the left POG, there are several experiments that support hyperactivation in the FL(Brewer *et al*., 2007). White matter disruptions have been reported as the main cause of functional abnormalities that arise with Schizophrenia (Cetin-Karayumak *et al*., 2019). The basal ganglia (BG) have been consistently investigated as a potential contributor to the development of psychosis in schizophrenia. This region contains a significant population of dopamine neurons and is therefore a key target for the development of anti-psychotic drugs (Perez-Costas, Melendez-Ferro and Roberts, 2010). Aberrations in the basal ganglia have been observed in a variety of ways across schizophrenic patients. Notably, numerous studies have reported a reduction in both bilateral caudate size and volume in these individuals (Ellison-Wright *et al*., 2008). In addition, altered white matter tract integration has been observed in the BG of schizophrenia patients when compared to healthy controls (Duan *et al*., 2015). The bilateral caudate is an essential component of planning and task execution, a function that is significantly impaired in individuals with schizophrenia. Collectively, these alterations lead to aberrant activation patterns in the SZ patients.

Previous studies on SZ suggest that aberrant activation in the third ventricle and right lateral ventricle may be related to deficits in emotional regulation, executive function, and white matter integrity in schizophrenia (Minzenberg *et al*., no date; Kubicki *et al*., 2007a; Li *et al*., 2007). Guo et al. reported reduced activation in the 3^rd^ ventricle in SZ patients during emotional processing tasks indicating deficits in emotional regulation in SZ patients (Guo, Liu, Xiao, *et al*., 2015). Mizenberg et al. found aberrant activation in the right lateral ventricle in individuals with SZ during tasks requiring executive function, which could be an indicator of suppressed cognitive processing in SZ patients (Minzenberg *et al*., no date). DTI studies have also shown structural abnormalities in 3^rd^ ventricle and right lateral ventricle, suggesting abnormalities in neural connectivity measured through fractional anisotropy (FA) in these regions (Kubicki *et al*., 2007b). The strong association between the size of the third and lateral ventricles in individuals with schizophrenia implies that there may be a causal interaction between the two structures. The expansion of both the third and lateral ventricles may be more indicative of schizophrenia than the enlargement of either structure alone (Kubicki *et al*., 2007b). Cerebellum has also shown to play a part in psychosis (Chang *et al*., 2022) and studies have been done in order to ascertain the mechanism behind this dysfunction (Kim *et al*., 2021). The reason why the activity for cerebellum is higher in patients than controls may be due to this dysfunction which misinterprets internal signal from auditory cortex as an external stimulus leading to hallucinations, a common symptom in schizophrenic patients (Bielawski and Bondurant, 2015).

The correlation matrices (**Figure 2)** show the voxel activation correlation amongst the ROIs and the results show an overall higher correlation for the controls. Previous research also supports these findings as patients with SZ have shown aberrant activation and altered connectivity between the ROIs (Duan *et al*., 2015b; Jiang *et al*., 2018; Zhao *et al*., 2018; Kraguljac *et al*., 2019; Bobilev, Perez and Tamminga, 2020) . This indicates that SZ increases the activation in all brain regions aberrantly leading to a lower correlation among them. It also supplements our findings of higher average voxel activations (**Figure 1**) in all ROIs.

In order to study the functional connectivity between the four ROIs in a time series, the study utilized vector auto-regression (VAR) model (**Figure 3**). VAR model shows hypo-connectivity in the patients supplementing previous research which also observed reduced degree centrality (DC) in patients with SZ compared to the controls. DC is an index used to identify the regions of the brain (at the whole-brain level) and display functional deficits in patients with SZ (Duan *et al*., 2017) leading to reduced functional connectivity. A lack of causal connectivity between the sensorimotor cortex (SMC) and multiple regions of the default mode network (DMN) in patients with SZ was also observed (Wertz *et al*., 2019), (Kottaram *et al*., 2019). Alterations to thalamic nuclei functional connectivity have also been reported in the cortico-cerebellar-thalamo-cortical circuit pathways of patients(Woodward, Karbasforoushan and Heckers, 2012) with SZ. GC was used to verify the above-mentioned results and elucidate the causal effects of the ROIs and the results confirmed lowered connectivity amongst the patients compared to the healthy controls. Previous research also supports our hypothesis stating that functional connectivity and causal relations decrease through the prognosis of SZ (Wismüller and Vosoughi, 2021). A study employing GC to identify and analyze the causal relations within the brain reported a decrease in the functional connectivity between the targeted areas of the brain (Koshiyama *et al*., 2021). Hence, GC can be used as a technique to create a causal map of the connections in healthy and diseased brain (Wismüller and Vosoughi, 2021). Lastly mediation analysis revealed a stronger partial mediation in the patients which could be attributed towards loss in connectivity. A similar pattern of altered rsFC has been observed throughout the entire brain in patients with SZ (Zhou *et al*., 2008)(Zhuo *et al*., 2018).

Amongst the strengths of this study, is its use of a combination of three different FC analysis methods to investigate functional disconnections between various regions of the brain. However, the study does have some limitations, which should be considered when interpreting the results. Most significantly, the sample size in the present study is small. Additional longitudinal follow-up studies with larger samples are needed to elucidate the alterations in FC between regions of the brain in patients with SZ. Furthermore, many studies have reported different results (Jiang *et al*., 2017) in the connectivity of various regions of the brain and more research needs to be done with subjects from several ethnicities so the differences can be recorded, and a standard diagnostic procedure can be implemented.

Together, these previous findings demonstrate that patients with SZ exhibit altered FC and effective connectivity (EC) among several brain regions.

## 5 Tables

**Table 1.** Average voxel activation values for the patients and controls.

| Average voxel activation |  |  |  |
| --- | --- | --- | --- |
| Lobe | Brain area | Activation in Patients | Activation in Controls |
| Frontal Lobe | Left middle frontal gyrus | 164.1021159 | 147.712822 |
|  | Right middle frontal gyrus | 201.8846313 | 190.551855 |
|  | Right superior frontal gyrus medial segment | 197.4187883 | 187.135653 |
|  | Left superior frontal gyrus medial segment | 189.5789453 | 178.038965 |
|  | Right superior frontal gyrus | 140.2225369 | 152.597001 |
|  | Right frontal operculum | 182.1029834 | 158.985179 |
|  | Right triangular part of the inferior frontal gyrus | 181.3794161 | 156.842199 |
|  | Right opercular part of the inferior gyrus | 146.5843352 | 130.320612 |
|  | Left opercular part of the inferior gyrus | 180.0882418 | 169.985293 |
|  | Left posterior orbital gyrus | 196.0969863 | 180.433215 |
| Basal ganglia | Right caudate | 194.3676695 | 179.890929 |
|  | Left accumbens area | 167.2095361 | 158.63256 |
|  | Right accumbens area | 177.7478272 | 162.647039 |
| Ventricular system | Right lateral ventricle | 170.582288 | 154.732335 |
|  | 3<sup>rd</sup> ventricle | 188.2889905 | 167.356182 |
| Cerebellum | Right cerebellum white matter | 165.826113 | 150.275748 |

**Table 2.** Correlation co-efficients for 16 regions of interest. 1. Left posterior orbital gyrus, 2. Right lateral ventricle, 3. 3^rd^ ventricle, 4. Right caudate, 5. Right cerebellum white matter, 6. Right accumbens area, 7. Left accumbens area, 8. Right triangular part of the inferior frontal gyrus, 9. Right superior frontal gyrus, 10. Left superior frontal gyrus medial segment, 11. Right superior frontal gyrus medial segment, 12. Right Frontal operculum, 13. Left middle frontal gyrus, 14. Right middle frontal gyrus, 15. Left opercular part of the inferior gyrus, 16. Right opercular part of the inferior gyrus. The upper half shows the controls, and the bottom half shows correlation values for the patients.

| 1 | 2 | 3 | 4 | 5 | 6 | 7 | 8 | 9 | 10 | 11 | 12 | 13 | 14 | 15 | 16 |
| --- | --- | --- | --- | --- | --- | --- | --- | --- | --- | --- | --- | --- | --- | --- | --- |
| 1.000 | -0.599 | -0.542 | -0.245 | 0.495 | 0.742 | 0.378 | 0.839 | 0.862 | 0.808 | 0.822 | 0.336 | 0.851 | 0.851 | 0.856 | 0.721 |
| -0.636 | 1.000 | 0.894 | 0.856 | -0.060 | -0.375 | 0.104 | -0.624 | -0.732 | -0.320 | -0.435 | 0.259 | -0.633 | -0.702 | -0.663 | -0.256 |
| -0.380 | 0.750 | 1.000 | 0.743 | -0.163 | -0.389 | 0.108 | -0.596 | -0.718 | -0.347 | -0.465 | 0.186 | -0.585 | -0.682 | -0.596 | -0.244 |
| -0.294 | 0.844 | 0.715 | 1.000 | 0.265 | 0.016 | 0.356 | -0.260 | -0.380 | 0.060 | -0.058 | 0.510 | -0.268 | -0.333 | -0.319 | 0.104 |
| 0.672 | -0.486 | -0.367 | -0.297 | 1.000 | 0.656 | 0.401 | 0.544 | 0.518 | 0.624 | 0.599 | 0.476 | 0.466 | 0.528 | 0.440 | 0.579 |
| 0.852 | -0.416 | -0.185 | -0.080 | 0.596 | 1.000 | 0.579 | 0.707 | 0.744 | 0.715 | 0.727 | 0.373 | 0.674 | 0.720 | 0.631 | 0.610 |
| 0.903 | -0.574 | -0.290 | -0.263 | 0.665 | 0.915 | 1.000 | 0.299 | 0.307 | 0.489 | 0.413 | 0.433 | 0.400 | 0.315 | 0.327 | 0.404 |
| 0.795 | -0.384 | -0.202 | -0.037 | 0.572 | 0.687 | 0.709 | 1.000 | 0.908 | 0.785 | 0.844 | 0.423 | 0.855 | 0.920 | 0.871 | 0.810 |
| 0.922 | -0.685 | -0.434 | -0.313 | 0.684 | 0.799 | 0.864 | 0.813 | 1.000 | 0.800 | 0.867 | 0.303 | 0.912 | 0.974 | 0.874 | 0.705 |
| 0.898 | -0.475 | -0.209 | -0.077 | 0.501 | 0.801 | 0.833 | 0.775 | 0.886 | 1.000 | 0.955 | 0.579 | 0.807 | 0.798 | 0.747 | 0.765 |
| 0.842 | -0.365 | -0.130 | 0.045 | 0.436 | 0.784 | 0.777 | 0.776 | 0.843 | 0.962 | 1.000 | 0.528 | 0.811 | 0.856 | 0.772 | 0.760 |
| -0.358 | 0.777 | 0.644 | 0.776 | -0.335 | -0.253 | -0.366 | 0.019 | -0.356 | -0.198 | -0.093 | 1.000 | 0.358 | 0.366 | 0.307 | 0.700 |
| 0.898 | -0.516 | -0.261 | -0.125 | 0.620 | 0.795 | 0.828 | 0.848 | 0.914 | 0.910 | 0.877 | -0.184 | 1.000 | 0.935 | 0.915 | 0.740 |
| 0.862 | -0.538 | -0.310 | -0.147 | 0.628 | 0.769 | 0.805 | 0.882 | 0.933 | 0.860 | 0.852 | -0.154 | 0.948 | 1.000 | 0.896 | 0.763 |
| 0.819 | -0.333 | -0.142 | -0.006 | 0.597 | 0.728 | 0.761 | 0.831 | 0.768 | 0.761 | 0.727 | 0.003 | 0.837 | 0.808 | 1.000 | 0.759 |
| 0.612 | -0.114 | 0.040 | 0.206 | 0.383 | 0.576 | 0.559 | 0.840 | 0.629 | 0.651 | 0.676 | 0.318 | 0.729 | 0.776 | 0.802 | 1.000 |

**Table 3(A).** Vector Autoregression (Average Controls). The table shows connectivity between 16 regions of interest. Right OpIFG: Right opercular part of the inferior gyrus, Right lat vent: Right lateral ventricle, Right CWM: Right cerebellum white matter, Left OpIFG: Left opercular part of the inferior gyrus, Right cau: Right caudate, Right TriFG: Right triangular part of the inferior frontal gyrus, 3^rd^ vent: 3^rd^ ventricle, Right MFG: Right middle frontal gyrus, Left acc: Left accumbens area, Left MSFG: Left superior frontal gyrus medial segment, Right MSFG: Right superior frontal gyrus medial segment, Left MFG: Left middle frontal gyrus, Right FO: Right frontal operculum, Left POrG: Left posterior orbital gyrus, Right acc: Right accumbens area, Right SFG: Right superior frontal gyrus, ROIs: Regions of interest, t: Time, Co-eff: Co-efficient, Std-error: Standard error, z: z-score

| ROIs | t-1 |  |  |  | t-2 |  |  |  |
| --- | --- | --- | --- | --- | --- | --- | --- | --- |
|  | Co-eff | Std-Error | z | p-value | Co-eff | Std-Error | z | p-value |
| 1. Right OpIFG←Right cau | -.0433 | .0871374 | -0.50 | 0.619 | .2054 | .0880396 | 2.33 | 0.020 |
| 2. Right OpIFG←Right lat vent | -.1890 | .0826249 | -2.29 | 0.022 | -.1887 | .0832408 | -2.27 | 0.023 |
| 3. Right OpIFG←3<sup>rd</sup> vent | -.0171 | .0303165 | -0.56 | 0.572 | .0902 | .0306975 | 2.94 | 0.003 |
| 4. Left OpIFG←Right cau | .0743 | .0921731 | 0.81 | 0.420 | .1923 | .0931274 | 2.07 | 0.039 |
| 5. Left OpIFG←Right lat vent | -.2386 | .0873998 | -2.73 | 0.006 | -.2023 | .0880513 | -2.30 | 0.022 |
| 6. Left OpIFG←Right acc | .0231 | .0538675 | 0.43 | 0.668 | -.1447 | .0533581 | -2.71 | 0.007 |
| 7. Left OpIFG←3<sup>rd</sup> vent | -.0170 | .0320685 | -0.53 | 0.595 | .0955 | .0324716 | 2.94 | 0.003 |
| 8. Right MFG←Right cau | -.0277 | .0850966 | -0.33 | 0.744 | .2312 | .0859777 | 2.69 | 0.007 |
| 9. Right MFG←Right lat vent | -.2474 | .0806897 | -3.07 | 0.002 | -.1618 | .0812913 | -1.99 | 0.046 |
| 10. Left MFG←Right cau | -.0965 | .0979631 | -0.99 | 0.324 | .3452 | .0989774 | 3.49 | 0.000 |
| 11. Left MFG←Right lat vent | -.2150 | .09289 | -2.31 | 0.021 | -.2546 | .0935825 | -2.72 | 0.007 |
| 12. Left MFG←Right acc | .0211 | .0572513 | 0.37 | 0.711 | -.1319 | .05671 | -2.33 | 0.020 |
| 13. Left MFG←3<sup>rd</sup> vent | -.0154 | .034083 | -0.45 | 0.650 | .0831 | .0345113 | 2.41 | 0.016 |
| 14. Left MFG←Left acc | -.0166 | .0536536 | -0.31 | 0.756 | .1172 | .0528586 | 2.22 | 0.027 |
| 15. Right cau←Right CWM | .1820 | .0713987 | 2.55 | 0.011 | .1777 | .0705207 | 2.52 | 0.012 |
| 16. Right cau←3<sup>rd</sup> vent | .0058 | .0327924 | 0.18 | 0.859 | .0823 | .0332045 | 2.48 | 0.013 |
| 17. Right MSFG←Right cau | .0448 | .1110214 | 0.40 | 0.686 | .2307 | .1121709 | 2.06 | 0.040 |
| 18. Right MSFG←Right MSFG | .4994 | .132555 | 3.77 | 0.000 | .2804 | .1351265 | 2.08 | 0.038 |
| 19. Right MSFG←Right CWM | .1124 | .0841007 | 1.34 | 0.181 | .2364 | .0830665 | 2.85 | 0.004 |
| 20. Right MSFG←3<sup>rd</sup> vent | -.0674 | .0386262 | -1.75 | 0.081 | .0783 | .0391116 | 2.00 | 0.045 |
| 21. Right MSFG←Left acc | -.0704 | .0608056 | -1.16 | 0.247 | .1328 | .0599045 | 2.22 | 0.027 |
| 22. Left MSFG←Right CWM | .1201 | .0901566 | 1.33 | 0.183 | .2006 | .089048 | 2.25 | 0.024 |
| 23. Left MSFG←3<sup>rd</sup> vent | -.0737 | .0414076 | -1.78 | 0.075 | .0946 | .041928 | 2.26 | 0.024 |
| 24. Left MSFG←Left acc | -.0728 | .0651841 | -1.12 | 0.264 | .1497 | .0642181 | 2.33 | 0.020 |
| 25. Right lat vent←Right acc | .0345 | .0553263 | 0.62 | 0.533 | -.1265 | .0548031 | -2.31 | 0.021 |
| 26. Right lat vent←Right TriFG | .0116 | .1023446 | 0.11 | 0.910 | -.2309 | .1027777 | -2.25 | 0.025 |
| 27. Right lat vent←3<sup>rd</sup> vent | .00069 | .032937 | 0.02 | 0.983 | .0955 | .0333509 | 2.86 | 0.004 |
| 28. Right lat vent←Left acc | -.0591 | .0518496 | -1.14 | 0.254 | .1689 | .0510812 | 3.31 | 0.001 |
| 29. Right SFG←Right lat vent | -.2580 | .0928791 | -2.78 | 0.005 | -.2237 | .0935715 | -2.39 | 0.017 |
| 30. Right SFG←Left Acc | -.0363 | .0536474 | -0.68 | 0.498 | .1319 | .0528524 | 2.50 | 0.013 |
| 31. Right acc←Right CWM | .1603 | .0941285 | 1.70 | 0.089 | .1886 | .0929711 | 2.03 | 0.042 |
| 32. Right TriFG←Right cau | -.0708 | .0786123 | -0.90 | 0.367 | .1565 | .0794262 | 1.97 | 0.049 |
| 33. Right TriFG←Right SFG | .0304 | .0880628 | 0.35 | 0.730 | -.1825 | .0889938 | -2.05 | 0.040 |

**Table 3(B).** Vector Autoregression (Average Patients). The table shows connectivity between 16 regions of interest in the patients. Right OpIFG: Right opercular part of the inferior gyrus, Right lat vent: Right lateral ventricle, Right CWM: Right cerebellum white matter, Left OpIFG: Left opercular part of the inferior gyrus, Right cau: Right caudate, Right TriFG: Right triangular part of the inferior frontal gyrus, 3^rd^ vent: 3^rd^ ventricle, Right MFG: Right middle frontal gyrus, Left acc: Left accumbens area, Left MSFG: Left superior frontal gyrus medial segment, Right MSFG: Right superior frontal gyrus medial segment, Left MFG: Left middle frontal gyrus, Right FO: Right frontal operculum, Left POrG: Left posterior orbital gyrus, Right acc: Right accumbens area, Right SFG: Right superior frontal gyrus, ROIs: Regions of interest, t: Time, Co-eff: Co-efficient, Std-error: Standard error, z: z-score

| ROIs | t-1 |  |  |  | t-2 |  |  |  |
| --- | --- | --- | --- | --- | --- | --- | --- | --- |
|  | Co-eff | Std-Error | z | p-value | Co-eff | Std-Error | z | p-value |
| 1.Right OpIFG←Right CWM | .0557 | .0300142 | 1.86 | 0.063 | .0841172 | .0306229 | 2.75 | 0.006 |
| 2.Right MFG←Right MFG | .5434 | .1209978 | 4.49 | 0.000 | .238683 | .1212015 | 1.97 | 0.049 |
| 3.Left MFG←Right SFG | -.3748 | .0848526 | -4.42 | 0.000 | .1731961 | .0877795 | 1.97 | 0.048 |
| **4.Right cau←Right cau** | .4165 | .097813 | 4.26 | 0.000 | .2204479 | .0962971 | 2.29 | 0.022 |
| **5.Right MSFG←Right MSFG** | .5197 | .1555654 | 3.34 | 0.001 | .4520271 | .1545717 | 2.92 | 0.003 |
| **6.Right MSFG←Right FO** | -.2769 | .0981718 | -2.82 | 0.005 | .2084026 | .0990462 | 2.10 | 0.035 |
| **7.Left MSFG←Right CWM** | .2036 | .0458128 | 4.44 | 0.000 | .0946744 | .0467419 | 2.03 | 0.043 |
| **8.Left MSFG←Right FO** | -.2680 | .1091011 | -2.46 | 0.014 | .2521826 | .1100729 | 2.29 | 0.022 |
| **9.Right lat vent←Right lat vent** | .0599 | .0805626 | 0.74 | 0.456 | .4517214 | .0805998 | 5.60 | 0.000 |
| **10.Right SFG←Right SFG** | .0511 | .0888479 | 0.58 | 0.564 | .2682422 | .0919125 | 2.92 | 0.004 |
| **11.Right SFG←Left acc** | .0239 | .0575091 | 0.42 | 0.677 | .1871581 | .0583278 | 3.21 | 0.001 |
| **12.Right acc←Left OpIFG** | -.0343 | .09703 | -0.35 | 0.724 | -<br>.2191728 | .0949006 | -<br>2.31 | 0.021 |
| **13.Right acc←Right lat vent** | -.4235 | .0933591 | -4.54 | 0.000 | .2103896 | .0934023 | 2.25 | 0.024 |
| **14.Right acc←Left acc** | .02267 | .0658204 | 0.34 | 0.730 | .2531285 | .0667575 | 3.79 | 0.000 |
| **15.Right TriFG←Right TriFG** | .5040 | .0953817 | 5.28 | 0.000 | .2058574 | .0944334 | 2.18 | 0.029 |
| **16.Right CWM←Right OpIFG** | -.3703 | .2052911 | -1.80 | 0.071 | .6117652 | .2058626 | 2.97 | 0.003 |
| **17.Right CWM←Right FO** | .5833 | .144118 | 4.05 | 0.000 | -<br>.3674006 | .1454016 | -<br>2.53 | 0.012 |
| **18.3<sup>rd</sup> vent←Right MSFG** | -.0599 | .2176579 | -0.28 | 0.783 | .4509556 | .2162676 | 2.09 | 0.037 |
| **19.3<sup>rd</sup> vent←3<sup>rd</sup> vent** | -.0298 | .0612107 | -0.49 | 0.626 | .2104276 | .0597504 | 3.52 | 0.000 |
| **20.3<sup>rd</sup> vent←Left acc** | -.1434 | .1051498 | -1.36 | 0.172 | .2434425 | .1066468 | 2.28 | 0.022 |
| **21.Left acc←Left acc** | .0615 | .0779635 | 0.79 | 0.430 | .4680959 | .0790735 | 5.92 | 0.000 |
| **22.Left POrG←3<sup>rd</sup> vent** | .0569 | .0341825 | 1.66 | 0.096 | -<br>.0662206 | .033367 | -<br>1.98 | 0.047 |
| **23.Left POrG←Left acc** | -.0166 | .0587198 | -0.25 | 0.803 | .1351963 | .0595557 | 2.27 | 0.023 |
| **24.Left POrG←Left POrG** | .3800 | .0823871 | 4.61 | 0.000 | .3183372 | .085313 | 3.73 | 0.000 |
| **25.Right FO←Right CWM** | .0572 | .0358747 | 1.60 | 0.110 | .1265024 | .0366023 | 3.46 | 0.001 |
| **26.Right FO←Right FO** | .3722 | .0854341 | 4.36 | 0.000 | .3410775 | .086195 | 3.96 | 0.000 |

**Table 4(A).**
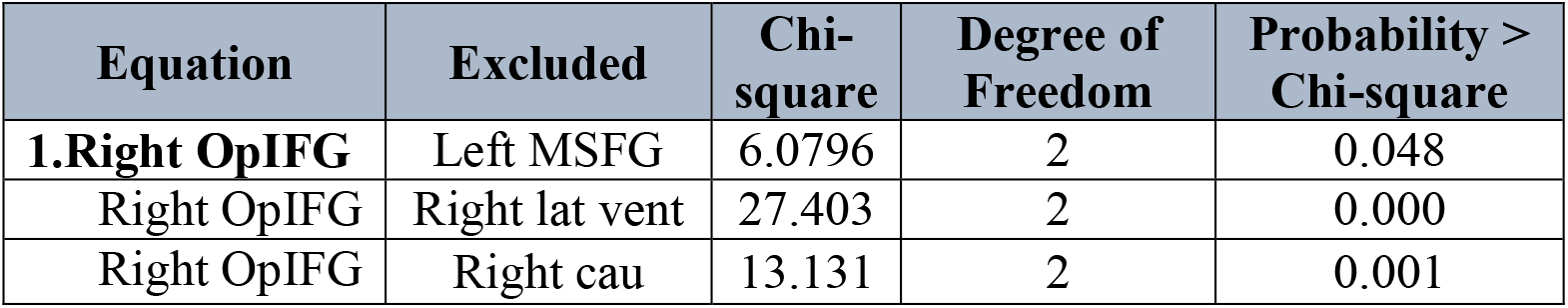
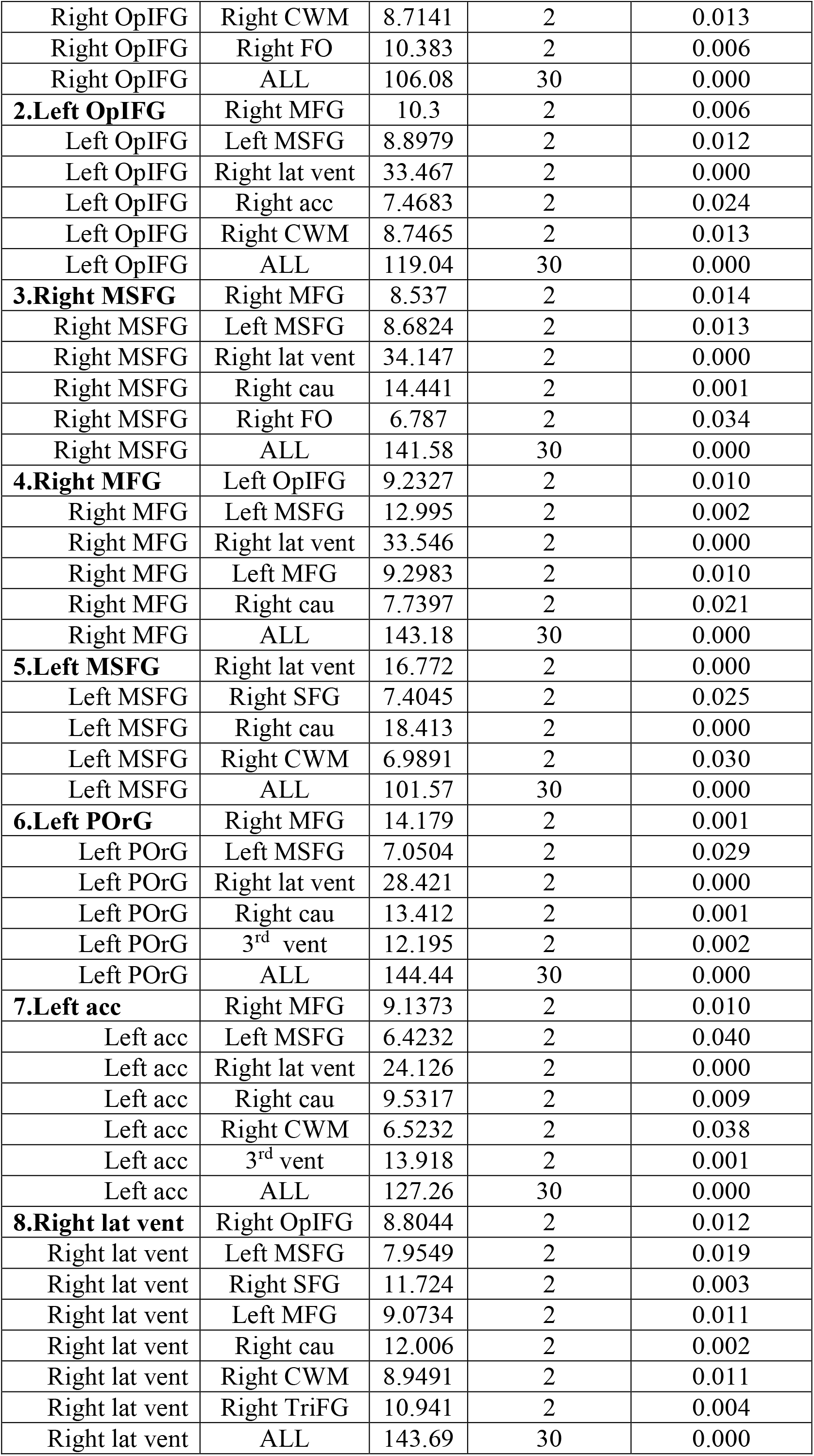
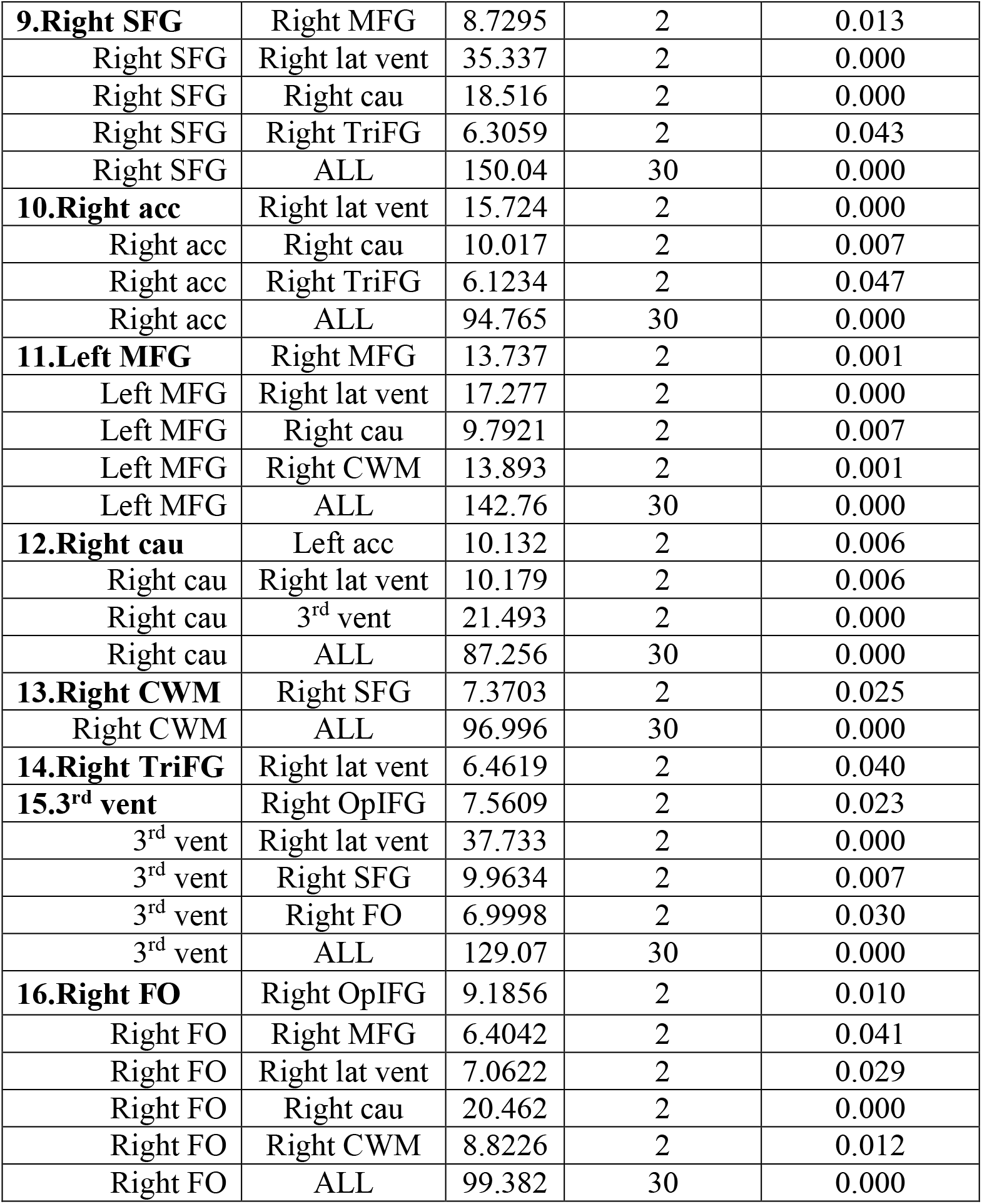
Granger Causality Test (Controls). The table shows intra-brain causal relations in the controls. Right OpIFG: Right opercular part of the inferior gyrus, Left OpIFG: Left opercular part of the inferior gyrus, Right MSFG: Right superior frontal gyrus medial segment, Right MFG: Right middle frontal gyrus, Left MSFG: Left superior frontal gyrus medial segment, Left POrG: Left posterior orbital gyrus, Left acc: Left accumbens area, Right Lat vent: Right lateral ventricle, Right SFG: Right superior frontal gyrus, Right acc: Right accumbens, Left MFG: Left middle frontal gyrus, Right cau: Right caudate, Right CWM: Right cerebellum white matter, Right TriFG: Right triangular part of the inferior frontal gyrus, 3^rd^ vent: 3^rd^ ventricle, Right FO: Right Frontal operculum, ALL: Causality

**Table 4(B).** Granger Causality Test (Patients). The table shows intra-brain causal relations in the patients. Right OpIFG: Right opercular part of the inferior gyrus, Left OpIFG: Left opercular part of the inferior gyrus, Right MSFG: Right superior frontal gyrus medial segment, Right MFG: Right middle frontal gyrus, Left MSFG: Left superior frontal gyrus medial segment, Left POrG: Left posterior orbital gyrus, Left acc: Left accumbens area, Right Lat vent: Right lateral ventricle, Right SFG: Right superior frontal gyrus, Right acc: Right accumbens, Left MFG: Left middle frontal gyrus, Right cau: Right caudate, Right CWM: Right cerebellum white matter, Right TriFG: Right triangular part of the inferior frontal gyrus, 3^rd^ vent: 3^rd^ ventricle, Right FO: Right Frontal operculum, ALL: Causality

| Equation | Excluded | Chi-square | Degree of Freedom | Probability > Chi-square |
| --- | --- | --- | --- | --- |
| **1.Right OpIFG** | Right cau | 13.123 | 2 | 0.001 |
| Right OpIFG | ALL | 77.53 | 30 | 0.000 |
| **2.Left OpIFG** | Right lat vent | 6.1263 | 2 | 0.047 |
| Left OpIFG | ALL | 76.164 | 30 | 0.000 |
| **3.Right MSFG** | Right lat vent | 6.1455 | 2 | 0.046 |
| Right MSFG | Right SFG | 13.451 | 2 | 0.001 |
| Right MSFG | Right cau | 20.511 | 2 | 0.000 |
| Right MSFG | ALL | 128.02 | 30 | 0.000 |
| **4.Right MFG** | Right lat vent | 6.4602 | 2 | 0.040 |
| Right MFG | Right SFG | 19.515 | 2 | 0.000 |
| Right MFG | Right cau | 15.408 | 2 | 0.000 |
| Right MFG | 3<sup>rd</sup> vent | 9.8246 | 2 | 0.007 |
| Right MFG | ALL | 128.15 | 30 | 0.000 |
| **5.Left MSFG** | Right lat vent | 7.3486 | 2 | 0.025 |
| Left MSFG | Right SFG | 17.802 | 2 | 0.000 |
| Left MSFG | Right cau | 12.036 | 2 | 0.002 |
| Left MSFG | ALL | 71.077 | 30 | 0.000 |
| **6.Left POrG** | Left acc | 7.7016 | 2 | 0.021 |
| Left POrG | Right lat vent | 6.4294 | 2 | 0.040 |
| Left POrG | Right SFG | 11.613 | 2 | 0.003 |
| Left POrG | Right cau | 19.438 | 2 | 0.000 |
| Left POrG | 3<sup>rd</sup> vent | 18.957 | 2 | 0.000 |
| Left POrG | Right FO | 8.3525 | 2 | 0.015 |
| Left POrG | ALL | 133.6 | 30 | 0.000 |
| **7.Left acc** | Left POrG | 7.6725 | 2 | 0.022 |
| Left acc | Right SFG | 13.988 | 2 | 0.001 |
| Left acc | Right cau | 27.731 | 2 | 0.000 |
| Left acc | 3<sup>rd</sup> vent | 20.377 | 2 | 0.000 |
| Left acc | Right FO | 7.2638 | 2 | 0.026 |
| Left acc | ALL | 162.23 | 30 | 0.000 |
| **8.Right lat vent** | Right SFG | 15.211 | 2 | 0.000 |
| Right lat vent | Right FO | 10.039 | 2 | 0.007 |
| Right lat vent | ALL | 65.785 | 30 | 0.000 |
| **9.Right SFG** | Right MSFG | 6.3137 | 2 | 0.043 |
| Right SFG | Left acc | 8.8369 | 2 | 0.012 |
| Right SFG | Right lat vent | 7.5767 | 2 | 0.023 |
| **10.Right SFG** | Right cau | 22.809 | 2 | 0.000 |
| Right SFG | Right TriFG | 10.825 | 2 | 0.004 |
| Right SFG | 3<sup>rd</sup> vent | 12.898 | 2 | 0.002 |
| Right SFG | ALL | 151.37 | 30 | 0.000 |
| **11.Right acc** | Left OpIFG | 8.5741 | 2 | 0.014 |
| Right acc | Left MSFG | 6.1715 | 2 | 0.046 |
| Right acc | Right lat vent | 20.748 | 2 | 0.000 |
| Right acc | Right cau | 6.2157 | 2 | 0.045 |
| Right acc | Right TriFG | 14.883 | 2 | 0.001 |
| Right acc | ALL | 108.39 | 30 | 0.000 |
| **12.Left MFG** | ALL | 83.196 | 30 | 0.000 |
| **13.Right cau** | Right OpIFG | 8.9789 | 2 | 0.011 |
| Right cau | Right SFG | 12.044 | 2 | 0.002 |
| Right cau | Right TriFG | 7.6104 | 2 | 0.022 |
| Right cau | Right FO | 16.49 | 2 | 0.000 |
| Right cau | ALL | 157.81 | 30 | 0.000 |
| **14.Right CWM** | Right SFG | 24.611 | 2 | 0.000 |
| Right CWM | Right cau | 11.155 | 2 | 0.004 |
| Right CWM | Right TriFG | 6.5353 | 2 | 0.038 |
| Right CWM | Right FO | 10.228 | 2 | 0.006 |
| Right CWM | ALL | 121.95 | 30 | 0.000 |
| **15.Right TriFG** | Right lat vent | 15.301 | 2 | 0.000 |
| Right TriFG | Right cau | 6.2818 | 2 | 0.043 |
| Right TriFG | 3<sup>rd</sup> vent | 8.5096 | 2 | 0.014 |
| Right TriFG | ALL | 106.38 | 30 | 0.000 |
| **16.3<sup>rd</sup> vent** | Right MFG | 8.2841 | 2 | 0.016 |
| 3<sup>rd</sup> vent | Right lat vent | 8.2368 | 2 | 0.016 |
| 3<sup>rd</sup> vent | Right CWM | 6.4066 | 2 | 0.041 |
| 3<sup>rd</sup> vent | ALL | 81.251 | 30 | 0.000 |
| **17.Right FO** | Left acc | 10.445 | 2 | 0.005 |
| Right FO | Right cau | 16.861 | 2 | 0.000 |
| Right FO | ALL | 70.743 | 30 | 0.000 |

**Table 5.** Mediation analysis for controls and patients. The table shows partial mediation in controls and patients. Left acc: Left accumbens area, Left MSFG: Left superior frontal gyrus medial segment, Right MSFG: Right superior frontal gyrus medial segment, Std-error: Standard error, * Significant.

|  | Type of Effect | Coefficient* | Std-Error | t-value | p-value | Mediation |
| --- | --- | --- | --- | --- | --- | --- |
| **Controls** | **1.Direct effect (Left acc on Left MSFG)** | 0.0691 | 0.0040 | 17.2897 | 0.0000 | **Partial Mediation** |
|  | **2.Direct effect (Right MSFG on Left MSFG)** | 0.8975 | 0.0135 | 66.5977 | 0.0000 |  |
|  | **3.Direct effect (Left acc on Right MSFG)** | -0.0743 | 0.0042 | -17.507 | 0.0000 |  |
|  | **4.Indirect effect (Left acc on Left MSFG)** | -0.0398 | - | - | - |  |
| **Patients** | **5.Direct effect (Left acc on Left MSFG)** | 0.1974 | 0.0173 | 11.3788 | 0.0000 | **Partial Mediation** |
|  | **6.Direct effect (Right MSFG on Left MSFG)** | 1.0227 | 0.0242 | 42.3007 | 0.0000 |  |
|  | **7.Direct effect (Left acc on Right MSFG)** | -0.0567 | 0.0174 | -3.2490 | 0.0013 |  |
|  | **8.Indirect effect (Left acc on Left MSFG)** | 0.5704 | - | - | - |  |

## 7 Declarations

### Ethics approval and consent to participate

#### IRB approval and number

The study was carried out in accordance with the code of international and local Ethics (Declaration of Helsinki). This study was reviewed and approved by the local ethics committee with approval no (NEU0334).

#### Patient consent for publication

The patient provided written informed consent for publication.

## 8 Abbreviations List

Right OpIFG: Right opercular part of the inferior gyrus
Right lat vent: Right lateral ventricle
Right CWM: Right cerebellum white matter
Left OpIFG: Left opercular part of the inferior gyrus
Right Cau: Right caudate
Right TriFG: Right triangular part of the inferior frontal gyrus
3^rd^ vent: 3^rd^ ventricle
Right MFG: Right middle frontal gyrus
Left acc: Left accumbens area
Left MSFG: Left superior frontal gyrus medial segment
Right MSFG: Right superior frontal gyrus medial segment
Left MFG: Left middle frontal gyrus
Right FO: Right frontal operculum
Left POrG: Left posterior orbital gyrus
Right acc: Right accumbens area
Right SFG: Right superior frontal gyrus
CWM: Cerebellum white matter
BG: Basal ganglia
VS: Ventricular system
FL: Frontal lobe
FC: Functional Connectivity

## 10 Author Contributions

### Study Design

Safee Ullah Chaudhary, Fayyaz Ahmad, Shahid Bashir, and Sadia Shakeel.

### Data Processing

Wajiha Abdullah, Alishba Tahir, Wjeeh ul Azeem, Mirza Ibrahim Ahmad, Muhammad Farhan Khalid, and Fayyaz Ahmed.

### Manuscript Writing

Wajiha Abdullah, Alishba Tahir, Wjeeh ul Azeem, Mirza Ibrahim Ahmad, Muhammad Farhan Khalid, Turki Abualait, Sadia Shakil, Shahid Bashir, Safee Ullah Chaudhary.

## 12 Acknowledgments

The authors would like to thank participants and their family members for their time and their data.

## Data Availability Statement

The datasets used and/or analyzed during the current study are available from the corresponding author on reasonable request.

## References

Abboud, R., Noronha, C. and Diwadkar, V.A. (2017) ‘Motor system dysfunction in the schizophrenia diathesis: Neural systems to neurotransmitters’, European Psychiatry, 44, pp. 125–133. Available at: 10.1016/j.eurpsy.2017.04.004.

Acar, E. et al. (2019) ‘Unraveling diagnostic biomarkers of schizophrenia through structure-revealing fusion of multi-modal neuroimaging data’, Frontiers in Neuroscience, 13(MAY), p. 416. Available at: 10.3389/FNINS.2019.00416/BIBTEX.

Algunaid, R.F. et al. (2018) ‘Schizophrenic patient identification using graph-theoretic features of resting-state fMRI data’, Biomedical Signal Processing and Control, 43, pp. 289–299. Available at: 10.1016/j.bspc.2018.02.018.

Andreasen, N.C. et al. (1995) ‘Correlational studies of the Scale for the Assessment of Negative Symptoms and the Scale for the Assessment of Positive Symptoms: an overview and update’, Psychopathology, 28(1), pp. 7–17.

Arnold, A., Liu, Y. and Abe, N. (2007) ‘Temporal causal modeling with graphical granger methods’, Proceedings of the ACM SIGKDD International Conference on Knowledge Discovery and Data Mining, pp. 66–75. Available at: 10.1145/1281192.1281203.

Beckmann, C.F. et al. (2005) ‘Investigations into resting-state connectivity using independent component analysis’, Philosophical Transactions of the Royal Society B: Biological Sciences, 360(1457), pp. 1001–1013. Available at: 10.1098/RSTB.2005.1634.

Bell, C.C. (1994) ‘DSM-IV: Diagnostic and Statistical Manual of Mental Disorders’, JAMA, 272(10), pp. 828–829. Available at: 10.1001/jama.1994.03520100096046.

Beveridge, N.J. and Cairns, M.J. (2012) ‘MicroRNA dysregulation in schizophrenia’, Neurobiology of disease, 46(2), pp. 263–271. Available at: 10.1016/J.NBD.2011.12.029.

Bielawski, M. and Bondurant, H. (2015) ‘Psychosis following a stroke to the cerebellum and midbrain: A case report’, Cerebellum and Ataxias, 2(1), pp. 1–4. Available at: 10.1186/S40673-015-0037-8/FIGURES/1.

Bighelli, I. et al. (2018) ‘Psychological interventions to reduce positive symptoms in schizophrenia: systematic review and network meta-analysis’, World psychiatry : official journal of the World Psychiatric Association (WPA), 17(3), pp. 316–329. Available at: 10.1002/WPS.20577.

Bobilev, A.M., Perez, J.M. and Tamminga, C.A. (2020) ‘Molecular alterations in the medial temporal lobe in schizophrenia’, Schizophrenia Research, 217, pp. 71–85. Available at: 10.1016/J.SCHRES.2019.06.001.

Brewer, W.J. et al. (2007) ‘Increased prefrontal cerebral blood flow in first-episode schizophrenia following treatment: longitudinal positron emission tomography study’, The Australian and New Zealand journal of psychiatry, 41(2), pp. 129–135. Available at: 10.1080/00048670601109899.

Buckley, P.F. and Miller, B.J. (2015) ‘Schizophrenia Research: A Progress Report’, The Psychiatric clinics of North America, 38(3), pp. 373–377. Available at: 10.1016/J.PSC.2015.05.001.

Calhoun, V.D. and Adali, T. (2012) ‘Multisubject independent component analysis of fMRI: A decade of intrinsic networks, default mode, and neurodiagnostic discovery’, IEEE Reviews in Biomedical Engineering, 5, pp. 60–73. Available at: 10.1109/RBME.2012.2211076.

Calhoun, V.D. and de Lacy, N. (2017) ‘Ten Key Observations on the Analysis of Resting-state Functional MR Imaging Data Using Independent Component Analysis’, Neuroimaging Clinics of North America, 27(4), pp. 561–579. Available at: 10.1016/j.nic.2017.06.012.

Cetin-Karayumak, S. et al. (2019) ‘White matter abnormalities across the lifespan of schizophrenia: a harmonized multi-site diffusion MRI study’, Molecular Psychiatry 2019 25:12, 25(12), pp. 3208– 3219. Available at: 10.1038/s41380-019-0509-y.

Chang, X. et al. (2022) ‘Alterations of cerebellar white matter integrity and associations with cognitive impairments in schizophrenia’, Frontiers in Psychiatry, 13. Available at: 10.3389/FPSYT.2022.993866/FULL.

Chatterjee, I. et al. (2020) ‘Impact of ageing on the brain regions of the schizophrenia patients: an fMRI study using evolutionary approach’, Multimedia Tools and Applications, 79(33–34), pp. 24757– 24779. Available at: 10.1007/S11042-020-09183-Z/FIGURES/6.

Chayer, C. and Freedman, M. (2001) ‘Frontal lobe functions’, Current Neurology and Neuroscience Reports 2001 1:6, 1(6), pp. 547–552. Available at: 10.1007/S11910-001-0060-4.

Chen, G. et al. (2011) ‘Vector autoregression, structural equation modeling, and their synthesis in neuroimaging data analysis’, Computers in Biology and Medicine, 41(12), pp. 1142–1155. Available at: 10.1016/J.COMPBIOMED.2011.09.004.

Christensen, J., Holcomb, J. and Garver, D.L. (2004) ‘State-related changes in cerebral white matter may underlie psychosis exacerbation’, Psychiatry Research: Neuroimaging, 130(1), pp. 71–78. Available at: 10.1016/J.PSCYCHRESNS.2003.08.002.

Corponi, F. et al. (2021) ‘Frontal lobes dysfunction across clinical clusters of acute schizophrenia’, Revista de Psiquiatría y Salud Mental [Preprint]. Available at: 10.1016/J.RPSM.2021.12.002.

Dimitrios Asteriou, S.G.H. (no date) ‘VAR Model’, in Applied econometrics. 3rd edn. Red Globe Press, p. 552.

Dong, D. et al. (2018) ‘Dysfunction of Large-Scale Brain Networks in Schizophrenia: A Meta-analysis of Resting-State Functional Connectivity’, Schizophrenia bulletin, 44(1), pp. 168–181. Available at: 10.1093/SCHBUL/SBX034.

Duan, M. et al. (2015) ‘Altered basal ganglia network integration in schizophrenia’, Frontiers in Human Neuroscience, 9(OCT), p. 561. Available at: 10.3389/fnhum.2015.00561.

Duan, M. et al. (2017) ‘[Degree centrality of the functional network in schizophrenia patients]’, Sheng wu yi xue gong cheng xue za zhi = Journal of biomedical engineering = Shengwu yixue gongchengxue zazhi, 34(6), pp. 837–841. Available at: 10.7507/1001-5515.201607062.

Ellison-Wright, I. et al. (2008) ‘The anatomy of first-episode and chronic schizophrenia: An anatomical likelihood estimation meta-analysis’, American Journal of Psychiatry, 165(8), pp. 1015–1023. Available at: 10.1176/APPI.AJP.2008.07101562/ASSET/IMAGES/LARGE/T416T3.JPEG.

Erdeniz, B. et al. (2017) ‘Decreased functional connectivity in schizophrenia: The relationship between social functioning, social cognition and graph theoretical network measures’, Psychiatry Research: Neuroimaging, 270, pp. 22–31. Available at: 10.1016/J.PSCYCHRESNS.2017.09.011.

Fan, Y.S. et al. (2021) ‘Individual-specific functional connectome biomarkers predict schizophrenia positive symptoms during adolescent brain maturation’, Human Brain Mapping, 42(5), pp. 1475–1484. Available at: 10.1002/HBM.25307.

Ghoshal, A. and Conn, P.J. (2015) ‘The hippocampo-prefrontal pathway: a possible therapeutic target for negative and cognitive symptoms of schizophrenia’, http://dx.doi.org/10.2217/fnl.14.63, 10(2), pp. 115–128. Available at: 10.2217/FNL.14.63.

Gillingham, S.M. et al. (2017) ‘Assessing cognitive functioning in ALS: A focus on frontal lobe processes’, Amyotrophic Lateral Sclerosis and Frontotemporal Degeneration, 18(3–4), pp. 182–192.

Gogtay, N. et al. (2011) ‘Age of Onset of Schizophrenia: Perspectives From Structural Neuroimaging Studies’, Schizophrenia Bulletin, 37(3), pp. 504–513. Available at: 10.1093/SCHBUL/SBR030.

Gohel, S. et al. (2018) ‘Frequency specific resting state functional abnormalities in psychosis’, Human Brain Mapping, 39(11), pp. 4509–4518. Available at: 10.1002/HBM.24302.

Grace, A.A. (2012) ‘Dopamine system dysregulation by the hippocampus: Implications for the pathophysiology and treatment of schizophrenia’, Neuropharmacology, 62(3), pp. 1342–1348. Available at: 10.1016/J.NEUROPHARM.2011.05.011.

Granger, C.W.J. (1969) ‘Investigating Causal Relations by Econometric Models and Cross-spectral Methods’, Econometrica, 37(3), p. 424. Available at: 10.2307/1912791.

Guo, W., Liu, F., Liu, J., et al. (2015) ‘Abnormal Causal Connectivity by Structural Deficits in First-Episode, Drug-Naive Schizophrenia at Rest’, Schizophrenia Bulletin, 41(1), pp. 57–65. Available at: 10.1093/SCHBUL/SBU126.

Guo, W., Liu, F., Xiao, C., et al. (2015) ‘Increased short-range and long-range functional connectivity in first-episode, medication-naive schizophrenia at rest’, Schizophrenia Research, 166(1–3), pp. 144– 150. Available at: 10.1016/J.SCHRES.2015.04.034.

Hare, S.M. et al. (2019) ‘Salience-Default Mode Functional Network Connectivity Linked to Positive and Negative Symptoms of Schizophrenia’, Schizophrenia bulletin, 45(4), pp. 892–901. Available at: 10.1093/SCHBUL/SBY112.

Hariri, A.R. (2019) ‘The Emerging Importance of the Cerebellum in Broad Risk for Psychopathology’, Neuron, 102(1), pp. 17–20. Available at: 10.1016/J.NEURON.2019.02.031.

Holt, D.J. et al. (2006) ‘Increased medial temporal lobe activation during the passive viewing of emotional and neutral facial expressions in schizophrenia’, Schizophrenia Research, 82(2–3), pp. 153– 162. Available at: 10.1016/J.SCHRES.2005.09.021.

Hua, M. et al. (2020) ‘Disrupted pathways from limbic areas to thalamus in schizophrenia highlighted by whole-brain resting-state effective connectivity analysis’, Progress in neuro-psychopharmacology & biological psychiatry, 99. Available at: 10.1016/J.PNPBP.2019.109837.

Huang, H. et al. (2018) ‘Altered corticostriatal pathway in first-episode paranoid schizophrenia: Resting-state functional and causal connectivity analyses’, Psychiatry Research: Neuroimaging, 272, pp. 38–45. Available at: 10.1016/J.PSCYCHRESNS.2017.08.003.

Huijgen, J. and Samson, S. (2015) ‘The hippocampus: A central node in a large-scale brain network for memory’, Revue Neurologique, 171(3), pp. 204–216. Available at: 10.1016/J.NEUROL.2015.01.557.

Jiang, Y. et al. (2017) ‘Common and distinct dysfunctional patterns contribute to triple network model in schizophrenia and depression: a preliminary study’, Progress in Neuro-Psychopharmacology and Biological Psychiatry, 79, pp. 302–310.

Jiang, Y. et al. (2018) ‘Progressive reduction in gray matter in patients with schizophrenia assessed with MR imaging by using causal network analysis’, Radiology, 287(2), pp. 633–642. Available at: 10.1148/RADIOL.2017171832/ASSET/IMAGES/LARGE/RADIOL.2017171832.FIG4.JPEG.

Johnsen, L.K. et al. (2020) ‘Alterations in Task-Related Brain Activation in Children, Adolescents and Young Adults at Familial High-Risk for Schizophrenia or Bipolar Disorder - A Systematic Review’, Frontiers in Psychiatry, 11, p. 632. Available at: 10.3389/FPSYT.2020.00632/BIBTEX.

Johnson, E.B. et al. (2015) ‘The impact of occipital lobe cortical thickness on cognitive task performance: An investigation in Huntington’s Disease’, Neuropsychologia, 79, pp. 138–146.

Kim, S.E. et al. (2021) ‘Impaired cerebro-cerebellar white matter connectivity and its associations with cognitive function in patients with schizophrenia’, npj Schizophrenia 2021 7:1, 7(1), pp. 1–7. Available at: 10.1038/s41537-021-00169-w.

Koenig, T. et al. (2001) ‘Decreased functional connectivity of EEG theta-frequency activity in first-episode, neuroleptic-naïve patients with schizophrenia: preliminary results’, Schizophrenia Research, 50(1–2), pp. 55–60. Available at: 10.1016/S0920-9964(00)00154-7.

Konradi, C. and Heckers, S. (2003) ‘Molecular aspects of glutamate dysregulation: implications for schizophrenia and its treatment’, Pharmacology & therapeutics, 97(2), p. 153. Available at: 10.1016/S0163-7258(02)00328-5.

Koshiyama, D. et al. (2020) ‘Neurophysiologic Characterization of Resting State Connectivity Abnormalities in Schizophrenia Patients’, Frontiers in Psychiatry, 11, p. 1350. Available at: 10.3389/FPSYT.2020.608154/BIBTEX.

Koshiyama, D. et al. (2021) ‘Neural network dynamics underlying gamma synchronization deficits in schizophrenia’, Progress in Neuro-Psychopharmacology and Biological Psychiatry, 107, p. 110224. Available at: 10.1016/J.PNPBP.2020.110224.

Kottaram, A. et al. (2019) ‘Brain network dynamics in schizophrenia: Reduced dynamism of the default mode network’, Human Brain Mapping, 40(7), pp. 2212–2228. Available at: 10.1002/HBM.24519.

Kubicki, M. et al. (2007a) ‘A review of diffusion tensor imaging studies in schizophrenia’, Journal of psychiatric research, 41(1–2), pp. 15–30. Available at: 10.1016/J.JPSYCHIRES.2005.05.005.

Kubicki, M. et al. (2007b) ‘A review of diffusion tensor imaging studies in schizophrenia’, Journal of psychiatric research, 41(1–2), pp. 15–30. Available at: 10.1016/J.JPSYCHIRES.2005.05.005.

Li, M. et al. (2011) ‘Genetic association and identification of a functional SNP at GSK3*β* for schizophrenia susceptibility’, Schizophrenia research, 133(1–3), pp. 165–171. Available at: 10.1016/J.SCHRES.2011.09.013.

Li, X. et al. (2007) ‘fMRI study of language activation in schizophrenia, schizoaffective disorder and in individuals genetically at high risk’, Schizophrenia research, 96(1–3), pp. 14–24. Available at: 10.1016/J.SCHRES.2007.07.013.

Lodge, D.J. and Grace, A.A. (2007) ‘Aberrant Hippocampal Activity Underlies the Dopamine Dysregulation in an Animal Model of Schizophrenia’, The Journal of Neuroscience, 27(42), pp. 11424 LP – 11430. Available at: 10.1523/JNEUROSCI.2847-07.2007.

Lodge, D.J. and Grace, A.A. (2011a) ‘Hippocampal dysregulation of dopamine system function and the pathophysiology of schizophrenia’, Trends in pharmacological sciences, 32(9), pp. 507–513. Available at: 10.1016/J.TIPS.2011.05.001.

Lodge, D.J. and Grace, A.A. (2011b) ‘Hippocampal dysregulation of dopamine system function and the pathophysiology of schizophrenia’, Trends in Pharmacological Sciences, 32(9), pp. 507–513. Available at: 10.1016/J.TIPS.2011.05.001.

Lynall, M.-E. et al. (2010) ‘Functional connectivity and brain networks in schizophrenia’, Journal of Neuroscience, 30(28), pp. 9477–9487.

Mannell, M. V. et al. (2010) ‘Resting state and task-induced deactivation: A methodological comparison in patients with schizophrenia and healthy controls’, Human Brain Mapping, 31(3), pp. 424–437. Available at: 10.1002/HBM.20876.

Minzenberg, M.J. et al. (no date) Meta-analysis of 41 Functional Neuroimaging Studies of Executive Function in Schizophrenia, jamanetwork.com. Available at: 10.1001/archgenpsychiatry.2009.91.

Pattabiraman, K. et al. (2021) ‘Hippocampal Hyperactivity Emerges During a Critical Period in Mice’, Biological Psychiatry, 89(9), pp. S209–S210. Available at: 10.1016/J.BIOPSYCH.2021.02.531.

Perez-Costas, E., Melendez-Ferro, M. and Roberts, R.C. (2010) ‘Basal ganglia pathology in schizophrenia: dopamine connections and anomalies’, Journal of Neurochemistry, 113(2), pp. 287– 302. Available at: 10.1111/J.1471-4159.2010.06604.X.

Psychiatry.org - DSM-5 Fact Sheets (no date). Available at: https://www.psychiatry.org/psychiatrists/practice/dsm/educational-resources/dsm-5-fact-sheets (Accessed: 28 February 2023).

del Re, E.C. et al. (2019) ‘Diffusion abnormalities in the corpus callosum in first episode schizophrenia: Associated with enlarged lateral ventricles and symptomatology’, Psychiatry Research, 277, pp. 45–51. Available at: 10.1016/J.PSYCHRES.2019.02.038.

Rubinov, M. and Bullmore, E. (2013) ‘Schizophrenia and abnormal brain network hubs’, Dialogues in Clinical Neuroscience, 15(3), p. 339. Available at: 10.31887/DCNS.2013.15.3/MRUBINOV.

Salisbury, D.F. et al. (2022) ‘Pathological resting-state executive and language system perfusion in first-episode psychosis’, NeuroImage: Clinical, 36, p. 103261. Available at: 10.1016/J.NICL.2022.103261.

Salvador, R. et al. (2010a) ‘Overall brain connectivity maps show cortico-subcortical abnormalities in schizophrenia’, Human Brain Mapping, 31(12), pp. 2003–2014. Available at: 10.1002/HBM.20993.

Salvador, R. et al. (2010b) ‘Overall brain connectivity maps show cortico-subcortical abnormalities in schizophrenia’, Human Brain Mapping, 31(12), pp. 2003–2014. Available at: 10.1002/HBM.20993.

Schizophrenia (no date). Available at: https://www.who.int/news-room/fact-sheets/detail/schizophrenia (Accessed: 28 February 2023).

Schneider, C.L. et al. (2019) ‘Survival of retinal ganglion cells after damage to the occipital lobe in humans is activity dependent’, Proceedings of the Royal Society B, 286(1897), p. 20182733.

Seth, A.K., Barrett, A.B. and Barnett, L. (2015) ‘Granger causality analysis in neuroscience and neuroimaging’, Journal of Neuroscience, 35(8), pp. 3293–3297. Available at: 10.1523/JNEUROSCI.4399-14.2015.

Sheffield, J.M. and Barch, D.M. (2016) ‘Cognition and resting-state functional connectivity in schizophrenia’, Neuroscience & Biobehavioral Reviews, 61, pp. 108–120. Available at: 10.1016/J.NEUBIOREV.2015.12.007.

Shinba, T. et al. (2004) ‘Near-infrared spectroscopy analysis of frontal lobe dysfunction in schizophrenia’, Biological Psychiatry, 55(2), pp. 154–164. Available at: 10.1016/S0006-3223(03)00547-X.

Silverstein, B.H., Bressler, S.L. and Diwadkar, V.A. (2016) ‘Inferring the dysconnection syndrome in schizophrenia: Interpretational considerations on methods for the network analyses of fMRI data’, Frontiers in Psychiatry, 7(AUG), p. 132. Available at: 10.3389/FPSYT.2016.00132/BIBTEX.

Skudlarski, P. et al. (2010) ‘Brain Connectivity Is Not Only Lower but Different in Schizophrenia: A Combined Anatomical and Functional Approach’, Biological Psychiatry, 68(1), pp. 61–69. Available at: 10.1016/J.BIOPSYCH.2010.03.035.

Snitz, B.E. et al. (2005) ‘Lateral and medial hypofrontality in first-episode schizophrenia: Functional activity in a medication-naive state and effects of short-term atypical antipsychotic treatment’, American Journal of Psychiatry, 162(12), pp. 2322–2329. Available at: 10.1176/appi.ajp.162.12.2322.

Spalthoff, R., Gaser, C. and Nenadić, I. (2018) ‘Altered gyrification in schizophrenia and its relation to other morphometric markers’, Schizophrenia research, 202, pp. 195–202.

SPM12 Software - Statistical Parametric Mapping (no date). Available at: https://www.fil.ion.ucl.ac.uk/spm/software/spm12/: (Accessed24 February 2023).

Squire, L.R., Stark, C.E.L. and Clark, R.E. (2004) ‘The medial temporal lobe’, Annual review of neuroscience, 27, pp. 279–306. Available at: 10.1146/ANNUREV.NEURO.27.070203.144130.

Subramaniam, M. et al. (2021) ‘Lifetime Prevalence and Correlates of Schizophrenia and Other Psychotic Disorders in Singapore’, Frontiers in psychiatry, 12. Available at: 10.3389/FPSYT.2021.650674.

Sutcliffe, G. et al. (2016) ‘Neuroimaging Intermediate Phenotypes of Executive Control Dysfunction in Schizophrenia’, Biological Psychiatry: Cognitive Neuroscience and Neuroimaging, 1(3), pp. 218– 229. Available at: 10.1016/J.BPSC.2016.03.002.

The epidemiology of early schizophrenia. Influence of age and gender on onset and early course - PubMed (no date). Available at: https://pubmed.ncbi.nlm.nih.gov/8037899/(Accessed: 28 February 2023).

Tordesillas-Gutierrez, D. et al. (2018) ‘The right occipital lobe and poor insight in first-episode psychosis’, Plos One, 13(6), p. e0197715.

Vanes, L.D. et al. (2019) ‘Neural correlates of positive and negative symptoms through the illness course: an fMRI study in early psychosis and chronic schizophrenia’, Scientific Reports 2019 9:1, 9(1), pp. 1–10. Available at: 10.1038/s41598-019-51023-0.

Venkataraman, A. et al. (2012) ‘Whole brain resting state functional connectivity abnormalities in schizophrenia’, Schizophrenia Research, 139(1–3), pp. 7–12. Available at: 10.1016/j.schres.2012.04.021.

Vos, T. et al. (2017) ‘Global, regional, and national incidence, prevalence, and years lived with disability for 328 diseases and injuries for 195 countries, 1990-2016: a systematic analysis for the Global Burden of Disease Study 2016’, Lancet (London, England), 390(10100), pp. 1211–1259. Available at: 10.1016/S0140-6736(17)32154-2.

Warburton, A. et al. (2016) ‘A GWAS SNP for Schizophrenia Is Linked to the Internal MIR137 Promoter and Supports Differential Allele-Specific Expression’, Schizophrenia bulletin, 42(4), pp. 1003–1008. Available at: 10.1093/SCHBUL/SBV144.

Wertz, C.J. et al. (2019) ‘Disconnected and Hyperactive: A Replication of Sensorimotor Cortex Abnormalities in Patients With Schizophrenia During Proactive Response Inhibition’, Schizophrenia Bulletin, 45(3), pp. 552–561. Available at: 10.1093/SCHBUL/SBY086.

Wildgust, H.J., Hodgson, R. and Beary, M. (2010) ‘The paradox of premature mortality in schizophrenia: new research questions.’, Journal of psychopharmacology (Oxford, England), 24(4 Suppl), pp. 9–15. Available at: 10.1177/1359786810382149.

Wismüller, A. and Vosoughi, M.A. (2021) ‘Classification of schizophrenia from functional MRI using large-scale extended Granger causality’, p. 49. Available at: 10.1117/12.2582039.

Woodward, N.D., Karbasforoushan, H. and Heckers, S. (2012) ‘Thalamocortical dysconnectivity in schizophrenia’, 169(10), pp. 1092–1099. Available at: https://ajp.psychiatryonline.org/doi/abs/10.1176/appi.ajp.2012.12010056 (Accessed: 14 February 2022).

Wu, D. and Jiang, T. (2020) ‘Schizophrenia-related abnormalities in the triple network: a meta-analysis of working memory studies’, Brain Imaging and Behavior, 14(4), pp. 971–980. Available at: 10.1007/S11682-019-00071-1/FIGURES/1.

Yang, C. et al. (2020) ‘Functional Alterations of White Matter in Chronic Never-Treated and Treated Schizophrenia Patients’, Journal of Magnetic Resonance Imaging, 52(3), pp. 752–763. Available at: 10.1002/JMRI.27028.

Yoon, J.H. et al. (2008) ‘Association of dorsolateral prefrontal cortex dysfunction with disrupted coordinated brain activity in schizophrenia: Relationship with impaired cognition, behavioral disorganization, and global function’, American Journal of Psychiatry, 165(8), pp. 1006–1014. Available at: 10.1176/appi.ajp.2008.07060945.

Zhang, M. et al. (2020) ‘Abnormal amygdala subregional-sensorimotor connectivity correlates with positive symptom in schizophrenia’, NeuroImage. Clinical, 26. Available at: 10.1016/J.NICL.2020.102218.

Zhou, B. et al. (2010) ‘Brain functional connectivity of functional magnetic resonance imaging of patients with early-onset schizophrenia’, Zhong nan da xue xue bao. Yi xue ban = Journal of Central South University. Medical sciences, 35(1), pp. 17–24. Available at: 10.3969/J.ISSN.1672-7347.2010.01.003.

Zhou, Y. et al. (2008) ‘Altered resting-state functional connectivity and anatomical connectivity of hippocampus in schizophrenia’, Schizophrenia Research, 100(1–3), pp. 120–132. Available at: 10.1016/J.SCHRES.2007.11.039.

Zhou, Z. et al. (2009) ‘Detecting directional influence in fMRI connectivity analysis using PCA based Granger causality’, Brain Research, 1289, pp. 22–29. Available at: 10.1016/J.BRAINRES.2009.06.096.

Zhuo, C. et al. (2018) ‘Altered resting-state functional connectivity of the cerebellum in schizophrenia’, Brain Imaging and Behavior, 12(2), pp. 383–389. Available at: 10.1007/S11682-017-9704-0/FIGURES/3.

Zovetti, N. et al. (2022) ‘Inefficient white matter activity in Schizophrenia evoked during intra and inter-hemispheric communication’, Translational Psychiatry, 12(1), pp. 1–11. Available at: 10.1038/s41398-022-02200-9.

